# RASAL3 regulates RAC/CDC42 GTPases, SAPK/JNK signaling, IL-2 gene activity, and directed motility in human T cells

**DOI:** 10.64898/2026.07.24.740537

**Authors:** Sabina Varadínková, Peter Oslacký, Štěpán Čada, Kamila Kvasničková, Petra Cigánková, Narendra V Gottumukkala, Burkhart Schraven, Jonathan A Lindquist, Michal Šmída

## Abstract

RASAL3 acts as a negative regulator of small cellular GTPases in hematopoietic cells. In immune cells, it primarily modulates the RAS/MAPK signaling pathway and affects cellular events including proliferation, differentiation, survival, and migration. Due to its inhibitory role in T cells, RASAL3 may represent a potential modulatory target for improving therapeutic strategies such as cell-based immunotherapy. However, most existing knowledge about RASAL3 function is derived from murine models, and its role in human T-cell signaling remains insufficiently characterized. To address this gap, we systematically investigated the function of RASAL3 in human primary T cells and T-cell line. For this purpose, we employed RASAL3 overexpression, CRISPR/Cas9-mediated deletion, and siRNA-mediated knockdown to thoroughly analyze the effects of RASAL3 on T-cell signaling, proliferation, and migration. Our data demonstrate that RASAL3 modulates primarily CDC42 and RAC1/RAC2 GTPases activity, SAPK/JNK phosphorylation, *c-Fos* and *c-Jun* expression, and IL-2 gene promoter activation. In addition, RASAL3 regulates actin polymerization and T-cell migration. Notably, loss of RASAL3 increases Jurkat T cells’ motility *in vivo* and potentiates their homing to the spleen. Collectively, these findings identify RASAL3 as an important regulator of human T-cell activation and motility and highlight its application potential for improving CAR-T cell therapy.

## INTRODUCTION

T lymphocytes play a central role in adaptive immunity by recognizing and responding to specific antigens. Stimulatory and inhibitory molecules tightly regulate T-cell activation and signaling to ensure effective immune responses while preventing autoimmunity (Sun et al. 2023). Among the key molecular regulators of T cells are small GTPases, which act as molecular switches controlling a wide range of cellular processes, including proliferation, differentiation, survival, gene expression, and cytoskeletal reorganization (Song et al. 2019). Two groups of proteins control GTPase activity with opposing action: guanine nucleotide exchange factors (GEFs) and GTPase-activating proteins (GAPs). GEFs activate small GTPases by facilitating the exchange of GDP for GTP, whereas GAPs inactivate them by accelerating GTP hydrolysis (Cherfils and Zeghouf 2013).

RASAL3 (RAS protein activator-like 3) belongs to the RAS-GAP protein family and contains PH (pleckstrin homology), C2, and RAS-GAP domains (Muro et al. 2015; Saito et al. 2015). RASAL3 is predominantly expressed by hematopoietic cells, and even though it was initially identified as a negative regulator of RAS-GTPase (Saito et al. 2015), it also seems to modulate RAC2-GTPase activity and thus partially influences AKT-mediated (AKT/protein kinase B) survival of hematopoietic cells (Shin et al. 2018a). Survival of murine naïve T and NK-T cells (natural killer) also depends on RASAL3, however it is dispensable for CD4+ or CD8+ T cellś development (Muro et al. 2015). Despite not affecting thymic or peripheral T cell development, the survival of naïve T cells upon RASAL3 loss in vitro appears unrelated to IL-7 stimulation, a crucial factor for T cell maturation (Hong et al. 2012), suggesting that *in vivo* response depends mainly on tonic TCR stimulation (Muro et al. 2018, 2015a). Moreover, RASAL3-deficient mice demonstrate a decreased number of NK-T cells in the liver, but not in the spleen or thymus. In addition, the liver damage induced by the α-galactosylceramide, a specific agonist for NK-T cells, is less severe in RASAL3 KO mice and is accompanied by decreased production of IL-4 and IFN-γ cytokines. In vitro, RASAL3-deficient NK-T cells also exhibit decreased IL-4 and IFN-γ secretion along with enhanced ERK phosphorylation (extracellular signal-regulated kinase), indicating that RASAL3 deficiency plays a pivotal role in regulating NK-T-mediated immune response (Saito et al. 2015). ERK phosphorylation and cell proliferation are also promoted by direct interaction of RASAL3 with the immune receptor CD229 in malignant multiple myeloma cells, where CD229 has recently emerged as a target for CAR-T cell therapy (chimeric antigen receptor) (Lin et al. 2022). Similarly, in cardiomyocytes, RASAL3 downregulation increases RAS/RAF/ERK signaling and improves cell survival under doxorubicin-induced cytotoxicity (Gao et al. 2023b), confirming its substantial role in regulating the RAS/MAPK (mitogen-activated protein kinase) signaling pathway.

Furthermore, RASAL3 expression, together with CD48 and P2RY10, positively correlates with improved overall survival in sarcoma patients, and this favorable prognosis is associated with enhanced CD8⁺ T cell infiltration (Zhu and Hou 2020). A similar pattern is observed in lung adenocarcinoma, where elevated RASAL3 expression corresponds to better prognosis and increased CD8⁺ T cell infiltration, although the underlying molecular mechanisms remain to be clarified (Liang et al. 2022). These findings suggest that RASAL3 not only regulates the development and activation of naïve T lymphocytes but may also play a critical role in controlling the migratory behavior of cytotoxic T cells and their recruitment into the tumor microenvironment.

In addition to its established roles in lymphocytes, RASAL3 also contributes to the regulation of myeloid cell function. Under inflammatory conditions, the RASAL3 promotes dendritic cell motility through modulating RHO-GTPase activity (Olivier et al. 2024). Furthermore, RASAL3 regulates the polarization of neutrophils and macrophages by orchestrating actin polymerization at the leading edge through mTORC2 signaling, thereby regulating actomyosin contractility, chemotaxis, and cell migration (Lin et al. 2024). Moreover, RASAL3 plays a pivotal role in regulating neutrophils’ immune response and preventing their hyperactivation in acute inflammatory conditions (Saito et al. 2021). Collectively, these findings indicate that RASAL3 acts as a broader regulator of cell motility across immune cell lineages, functioning not only in tumor-associated contexts but also in inflammatory environments.

Although previous studies collectively suggest that RASAL3 participates in the functional regulation of multiple immune cell types, most of these studies have been conducted in mice, and consequently, the role of RASAL3 in human T-cell signaling remains largely unexplored. Given that RASAL3 presumably acts as a negative regulator of RAS signaling (Saito et al. 2015a), we hypothesized that RASAL3 deficiency could amplify key T-cell signaling pathways and thereby potentiate T-cell physiological responses. Enhanced T-cell functionality is particularly relevant in the context of cellular immunotherapy, for example, CAR-T cell therapy, where improving the signaling capacity, persistence, and migratory potential of engineered T cells remains a major challenge (Liu et al. 2024).

To address this knowledge gap, we systematically investigated the role of RASAL3 in human T-cell signaling and migration, two key properties underlying T-cell functionality. Therefore, we modulated RASAL3 expression levels in a T-cell line by transient overexpression or by CRISPR/Cas9-guided knockout of RASAL3. We analyzed the impact of RASAL3 overexpression or deficiency on the activation of small GTPases, including RAS, RAC1, RAC2, RHO, and on major signaling pathways such as AKT, ERK, SAPK/JNK (stress-activated protein kinase/c-Jun N-terminal kinase), as well as the *c-Fos* and *c-Jun* expression. We also explored the consequences on the IL-2 gene promoter activation and cell proliferation. In addition, we assessed the effects of RASAL3 expression changes on cytoskeletal dynamics, cell adhesion, and migratory behavior. We also validated the impact of RASAL3 loss on cell migration *in vivo*. Importantly, we confirmed the key findings in human primary T cells.

Taken together, these analyses provide new insights into the role of RASAL3 in human T-cell signaling, activation, cytoskeleton reorganization, and migration.

## MATERIAL and METHODS

### Cells

Human Jurkat T-cell line (derived from acute T-cell lymphoblastic leukemia; obtained from DSMZ) was cultured in RPMI-1640 medium supplemented with 10 % fetal bovine serum (FBS) and 1 % penicillin–streptomycin (all reagents from Sigma-Aldrich). Cells were maintained at 37°C in a humidified atmosphere with 5 % CO₂. Cells were passaged every 2–3 days and cryopreserved/thawed according to standard protocols.

Primary human T cells were isolated from healthy human donors’ fresh blood using the Ficoll-Paque PLUS (Cytiva) gradient centrifugation. T cells were purified by non-T cell depletion using the Pan T cell Isolation Kit II (Miltenyi Biotec). T cells were cultured in RPMI-1640 medium (Gibco) supplemented with 10% FBS and 1% penicillin-streptomycin.

### Primary T-cells activation

A 24-well flat-bottom tissue culture plate (TPP) was pre-coated with 10 µg/mL rabbit-anti-mouse immunoglobulin (Dako) dissolved in phosphate-buffered saline (PBS) overnight at 4°C. After three washings with PBS, 1 µg/mL of the specific anti-CD3, anti-CD4, anti-CD28, and anti-ICAM antibodies (all Exbio) were immobilized on plates at 4°C overnight. After PBS washing, T cells (6 x 10^5^/mL) were inoculated in 1 mL complete RPMI per well. In parallel, T cells were inoculated into uncoated wells and stimulated with soluble stimulants: 10 nM phorbol ester PMA (Merck), 0.25 µg/mL Ionomycin (Merck), 10 µg/ml phytohemagglutinin PHA, 10 U/mL IL-2 (Tebu-Bio), 10 U/ml IL-10, and 200 ng/mL SDF-1α (PeproTech). Resting cells were cultured without any stimulus. After 3 days of incubation at 37°C and 5% CO_2_, the cells were harvested for analysis.

### RASAL3 cloning

Human peripheral blood T cells were cultured for 3 days on anti-CD3-pre-coated plastic plates. Then, total RNA was isolated using the Absolutely RNA Miniprep Kit (Agilent Technologies) according to the manufacturer’s instructions. The full-length RASAL3 cDNA was synthesized and amplified with the SuperScript One-Step RT-PCR System for Long Templates (Invitrogen) using gene-specific primers. The amplified product was inserted into the linearized pCR 2.1 vector provided with the TA Cloning Kit (Invitrogen). The construct was transformed into chemically competent E. coli, and positive colonies were identified by blue/white selection on plates containing ampicillin and X-Gal. Plasmid DNA was extracted from the positive clones and tested by restriction enzyme digestion for proper insert orientation and by sequencing for correct nucleotide sequence. Finally, the insert was cut out using the restriction sites within the polylinker sequence of pCR 2.1 vector and inserted into the expression vector pEF IRES.

### Cell transfection

Jurkat T cells (20 × 10^6^) were washed and resuspended in 350 µL PBS. Cells were electroporated with 30 µg plasmid DNA in a 4 mm cuvette using the Gene Pulser II (210 V, 950 µF) (Bio-Rad). Immediately after electroporation, 450 µL RPMI-1640 medium was added, the precipitated DNA removed, and cells were cultured in 40 mL complete RPMI-1640 medium for 24–72 h.

Human primary T cells (8 x 10^6^) were washed with PBS, resuspended in 200 µL Opti-MEM (Invitrogen), and transfected with 1.2 nmol pool of 4 siRNAs targeting RASAL3 or scrambled control siRNA (both Dharmacon) in a 4 mm cuvette with a square-wave pulse (1000 V, 0.5 ms, 2 pulses, gap between pulses 5 s) using the Gene Pulser Xcell Electroporation System (Bio-Rad).

### Immunofluorescence

Jurkat cells (7 x 10^6^/ml) transfected with the RASAL3 construct were pipetted into 20 µL RPMI-1640 media per spot on a 12-spot slide (Paul Marienfeld GmbH & Co. KG) and incubated at 37°C for 30 min in a humid chamber. Cells were washed, fixed with 1 % paraformaldehyde/ 0.05 % glutaraldehyde in PBS, and permeabilized using 0.1 % Triton X-100 in PBS for 10 min each. The slide was blocked with 1 % bovine serum albumin (BSA)/PBS, and cells were stained with the primary anti-RASAL3 antibody for 60 min in a humid chamber. After washing and additional blocking, the cells were stained with FITC-labeled secondary antibody in a humid chamber. Following additional washing, samples were mounted with Vectashield Mounting Media (Vector Laboratories Inc.). Confocal images were acquired using a Leica DMIRE2 microscope and processed with the LSM Image Browser software (Zeiss).

### Membrane fractionation

Jurkat or primary human T cells (40 x 10^6^) were washed with PBS, and cell pellets were snap-frozen in liquid nitrogen. The pellets were then lysed in ice-cold hypotonic lysis buffer (20 mM HEPES, 5 mM EDTA, 1 mM MgCl_2_, 1 mM PMSF, 5 mM sodium pyrophosphate, 1 mM sodium vanadate) and resuspended 15x times with a syringe. After 10 min incubation on ice, lysates were centrifuged at 100.000 g for 60 min at 4°C and the supernatant was removed as the cytosolic fraction. The pellets were resuspended in a lysis buffer (20 mM Tris, 1 % Triton X-100, 150 mM NaCl, 1 mM MgCl_2_, 1 mM PMSF, 1 mM sodium vanadate), kept on ice for 10 min, and centrifuged at 100.000 g, 60 min, 4°C. Supernatants containing the solubilized membrane proteins were preserved for Western blot analysis.

### TCR stimulation

Cells (2 × 10⁶) were starved for 1 hour in RPMI-1640 medium (Sigma-Aldrich) without serum, supplemented with 2 % BSA and 0.5 % HEPES (both Sigma-Aldrich). TCR signaling was induced by stimulating cells with 1 µL of anti-CD3 (MEM92) and 1 µL of anti-CD28 antibodies (CD28.2; both from Exbio) per 100 µL of cell suspension for the indicated time points. Reactions were stopped by adding 1 mL of ice-cold 1× Tris-buffered saline (TBS). Cells were then either lysed in RIPA buffer containing protease (PMSF) and phosphatase (PhosSTOP, both Roche) inhibitors for Western blot analysis, or collected in RLT buffer (RNeasy Mini Kit; Qiagen) for RNA extraction.

### Immunoprecipitation

Cells (20 x 10^6^) were washed in PBS and resuspended in 500 µL RIPA lysis buffer. An aliquot (10 %) was kept as a whole lysate, and the rest was incubated with 1 mg/mL BSA, anti-RASAL3 antibody or IgG control, and protein A Sepharose for 2 to 18 hours with rotation at 4°C. Beads were washed 5 times with low-detergent washing buffer (0.05 % NP-40, 5 mM EDTA, 150 mM NaCl, and 50 mM Tris, pH 7.4), and the immunoprecipitated material was released by cooking in reducing sample buffer.

### Western blotting

Cells (2 × 10⁶) were washed with PBS and lysed in RIPA buffer containing protease (PMSF) and phosphatase (PhosSTOP) inhibitors (both Roche), then boiled at 98 °C for 10 min. Proteins were separated by SDS-PAGE and blotted onto nitrocellulose or PVDF membranes. Membranes were blocked with 5 % BSA for 2 h at room temperature (RT) with constant rolling and incubated with primary antibodies diluted in 5 % BSA or non-fat dry milk according to the manufacturer’s instructions overnight at 4 °C on a roller. List of used primary antibodies is available in the Supplementary Fig. 3A. After three washes with TBS-Tween (TBST), membranes were incubated with HRP-conjugated secondary antibodies (anti-mouse or anti-rabbit, 1:10,000, Cell Signaling Technology) for 1 h at RT with constant rolling, washed three times with TBST, and developed with ECL (Bio-Rad). Chemiluminescence was detected using the Alliance Q9 imaging system (Uvitec Cambridge).

### Calcium measurement

Jurkat T-cells were transfected with either empty vector or RASAL3 construct together with a vector containing GFP. 24 hrs after transfection, 2 x 10^6^ cells were washed in RPMI-1640 medium without phenol red and loaded with Indo-1/AM (Invitrogen) for 45 min at 37°C. Cells were briefly washed and incubated for an additional 30 min before measuring the FL4 (510/20nm) vs. FL5 (400/40nm) ratio on a flow cytometer (BD LSR), while gating on GFP-positive cell population. During the measurement, cells were stimulated first with soluble anti-CD3 antibody (C305, 1:50) and then with ionomycin (2 µg/mL).

### Luciferase assay

Jurkat cells were electroporated with 20 µg RASAL3 plasmid plus 5 µg Firefly Luciferase under the complete IL-2 promoter and 1 µg Renilla luciferase constructs, as mentioned above. After 24 hours, cells were inoculated in quadruplicates onto 96-well plates (70.000 cells in 100 µL per well) pre-coated with anti-CD3, anti-CD28 and anti-CD3/CD28 antibodies. Control wells remained uncoated and the control cells received PBS (negative control) or the combination of 10 mM PMA plus 1 µg/mL ionomycin (positive control). After 6 hours at 37°C, cells were washed twice with PBS and lysed in 30 µL lysis buffer for 15 min at RT. The dual luminescence was detected using dual-luciferase reporter assay (Promega) on a single-injector luminometer. The Firefly luminescence signal was normalized to Renilla.

### Generation of RASAL3-knockout cells using CRISPR/Cas9

RASAL3-deficient Jurkat cells were generated by electroporation using the Neon™ Transfection System (Invitrogen). Briefly, cells (2 × 10⁵) were washed with PBS and resuspended in 5 µL of Buffer R per sample. A ribonucleoprotein (RNP) complex was prepared by mixing three single guide RNAs (sgRNAs, 30 µM, 3 µL/sample; Synthego; sequences listed in Supplementary Fig. 1C) with Cas9 protein (20 µM, 0.5 µL/sample; Synthego) in 3.5 µL of Buffer R. The mixture was incubated at RT for 10 min to allow RNP complex formation. The RNP complex, together with a GFP plasmid (0.5 µL/sample), was electroporated into Jurkat cells at 1325 V, 10 ms, 3 pulses. Cells were subsequently cultured for 2 days in RPMI-1640 medium without antibiotics. Single-cell sorting into 96-well plates was performed using flow cytometry (BD Biosciences) to establish individual RASAL3 knockout clones, which were expanded for 3 weeks. Editing efficiency was confirmed by genomic DNA sequencing (data not shown) and Western blot analysis (Supplementary Fig. 1A).

### Quantitative real-time PCR

For RNA expression analysis, cells were lysed in RLT lysis buffer supplemented with 2–mercaptoethanol (Sigma-Aldrich) and RNA was extracted using the RNeasy Mini Kit (Qiagen) according to the manufactureŕs protocol. Total RNA concentration and purity were verified using a NanoDrop 2000c (Thermo Fisher). 400 ng RNA was diluted in 11 µL of Nuclease-free (NF) water and incubated with 1 µL of Random Hexamer Primer (Thermo Scientific) for 5 min at 65 °C in a C1000 Touch TM Thermal Cycler (Bio-Rad). A mixture of 4 μL 5x Reaction Buffer, 2 μL dNTP mix, 1 μL RevertAid Reverse Transcriptase, and 0.5 μL Ribolock RNase Inhibitor (all Thermo Scientific) was added to each sample, and a Reverse Transcription was performed in a C1000 Touch TM Thermal Cycler (10 min/25°C, 60 min/45°C, 10 min/70°C). cDNA was further diluted in NF-water to a final concentration of 100 ng/µL per sample. Primers for *c-Fos*, *c-Jun*, Hprt, and Gapdh genes (all Generi Biotech; sequences listed in Supplementary Fig. 1B) were diluted in NF-water in a 1:1:8 ratio (Forward primer: Reverse primer: NF-water). 4 µL cDNA was mixed with 1 µL diluted primers and 5 μL PowerUP SYBR Green MasterMix (Applied Biosystems) and qPCR run using the QuantStudio 12 Flex Real-Time PCR system (Applied Biosystems). Hprt and Gapdh were used as housekeeping controls.

### Active GTPase pull-down assay

Cells (1 × 10⁷) were starved for 1 hour in serum-free RPMI-1640 medium containing 0.5 % BSA at 37 °C and 5 % CO₂. After washing, cells were stimulated with anti-CD3 (MEM92) and anti-CD28 (both Exbio) antibodies for the indicated time points. Stimulation was stopped with 1 mL ice-cold TBS, and cells were immediately lysed. Active GTPases were isolated using commercial pull-down kits according to the manufacturer’s instructions (Active CDC42, RAS, RAC1/RAC2, RAP1, and RHO Pull-Down and Detection Kits, Thermo Fisher Scientific). Active CDC42, RAS, RAC1, RAC2, RAP1, and RHO pull-downs, their total proteins, and loading controls were analyzed by Western blotting.

### Flow cytometry

Cells were washed with PBS and stained with 1 μL anti-CD3-FITC-conjugated antibody diluted in 50 μL PBS for 30 min on ice in the dark. After washing with PBS, fluorescence was analyzed by FACSVerse flow cytometer (BD Bioscience).

### Growth curve

Jurkat cells (3 × 10⁵) were seeded onto 12-well plates in triplicate. Viable cells were assessed by Trypan blue exclusion (Avantor Science Central), counted every 2–3 days using a Luna-II cell counter (Logos Biosystems), and reseeded at the same density. After three weeks, growth curves were calculated assuming exponential growth and expressed as the negative logarithm of the initial cell number.

### Cell cycle analysis

Jurkat cells were collected at three distinct time points, washed three times with PBS, and fixed with ice-cold 70 % ethanol for 2h at -20 °C. After washing with cold PBS, cells were incubated for 30 min in Vindel’s solution containing RNase A and propidium iodide (PI). Cell cycle distribution was analyzed by flow cytometry.

### Actin polymerization assay

Cells (2 × 10⁵) were starved for 1 h in serum-free RPMI-1640 medium, washed with PBS, and resuspended in 100 μL PBS per sample. Cells were stimulated with anti-CD3 (MEM92) and anti-CD28 antibodies (Exbio; 1 μL each) for indicated times at 37 °C with shaking (300 rpm), including unstimulated controls that received PBS. Reactions were stopped by adding 100 μL FA-T (3.7 % formaldehyde, 1 % Triton X-100 in PBS). TRITC–phalloidin (Thermo Fisher Scientific; 3 μL) was added, and samples were incubated for 15 min at RT in the dark, washed with PBS, and analyzed by flow cytometry.

### Migration assay

Cell migration was assessed using transwell inserts with 8 μm pore size for Jurkat cells and 5 μm pore size for primary T cells (Corning Costar). The lower chamber was filled with 600 μL migration media (RPMI-1640 supplemented with 2 % BSA and 2 % HEPES) with SDF-1α chemokine, CCL19, CCL21 (200 ng/mL; all PeproTech) or PBS in control wells. Cells (5 × 10⁵) were seeded onto inserts and allowed to migrate for 5 h at 37 °C in 5 % CO₂. Cells in the lower chamber were then collected and quantified by flow cytometry. Migrated cells were expressed as a percentage of the total cellś input.

### Adhesion assay

The 96–well plate was pre-coated with 10 µg/mL Fibronectin (Sigma-Aldrich) diluted in 2 % BSA/PBS at 4 °C overnight. Wells were washed three times with PBS without Ca²⁺/Mg²⁺, blocked for 2 h with 2 % BSA/PBS at 37°C and 5 % CO₂, and washed with Hank’s balanced salt solution (HBSS; Sigma-Aldrich). Jurkat cells were starved for 1 h in serum-free RPMI-1640 medium containing 0.5 % BSA, washed with HBSS, and pre-incubated for 15 min at 37 °C with SDF-1α (200 ng/mL; PeproTech), MnCl₂ (2 mM), anti-CD3(MEM92)/CD28.2 antibodies (10 μL/mL; Exbio), or PBS. Cells were seeded at 1 × 10⁵ cells/well in triplicate and allowed to adhere for 30 minutes. Non-adherent cells were removed by washing three times with HBSS. CellTiter-Glo (Promega; diluted 1:6 in PBS) was added (50 μL/well), plates were shaken for 20 min in the dark, and luminescence was measured using a Tecan plate reader.

### Time-lapse microscopy

Time-lapse microscopy was performed as described previously (Čada et al. 2022). Multi-melting agarose (Thermo Fisher Scientific) was dissolved in Milli-Q water and mixed with RPMI-1640 medium supplemented with 2× FBS and 2× HBSS. SDF-1α chemokine (Peprotech) was added to half of the mixture, which was then pipetted into 8-well plates (ibidi GmbH). After gel solidification, a small well was created, and Hoechst-stained cells were added beneath the agarose. Samples were imaged for 90 min using CellSens Dimension software (Olympus LS), and cell movement was analyzed with ImageJ Fiji.

### *in vivo* cell homing assay

NOD-scid IL2Rg^null^ mice (NSG; Charles River Laboratories, France) were housed in the animal facility of Masaryk University, Brno. All animal experiments were performed with the approval of the Masaryk University’s ethics committee and in accordance with the Act on Veterinary Care (Act No.⍰166/1999 Coll.) and the Act on the Protection of Animals Against Cruelty (Act No.⍰246/1992 Coll.).

RASAL3 KO or WT cells, alternatively, (2 × 10⁶ of each per mouse) were labeled with CellTracker CMTPX (200 nM; Invitrogen) and mixed with the unlabeled cell counterpart in a 1:1 ratio. Different target cells were stained in each replicate to exclude any effect of the staining itself on the cell motility. Mixed cells were intraperitoneally injected into three mice and allowed to migrate for 48 hours. Mice were sacrificed, and blood, bone marrow, spleen, and liver were collected. Tissues were homogenized, filtered, and cells washed with PBS. Erythrocytes were lysed using Erythrolysis buffer (1:3 ratio, 15 min at RT). Samples were stained with anti-CD3 antibody (FITC), and the proportion of KO versus WT cells was analyzed by flow cytometry.

### Viability assay

Cells (1 × 10⁵ cells/well) were seeded in 96-well plates in 100 µl of complete RMPI-1640 media in triplicate. Doxorubicin (1 mM) was 10 × serially diluted in DMSO (Serva GmbH) using 2-fold (clone 1) or 2.5-fold (clone 2) dilutions, and 1 μL of the final solution was added to each well. Cells were cultured for 3 days, then 20 μL of CellTiter-Glo (Promega) was added per well. Plates were shaken for 20 min, and luminescence, corresponding to cellular ATP levels, was measured using a Tecan plate reader.

### Cytokine production assay

Primary human T cells electroporated with RASAL3 siRNAs or scrambled siRNA were inoculated onto 96-well plates pre-coated with anti-CD3/CD28 antibodies and incubated at 37°C for 24 and 72 hours. Cytokine production was determined by LEGENDplex (8-plex) multiplex assay (Biolegend) according to the manufacturer’s instructions and measured on FACSCalibur (BD Bioscience).

### CFSE dilution assay

Primary human T cells electroporated with RASAL3 siRNAs or scrambled siRNA were washed once with PBS and loaded with 0.1 µM carboxyfluorescein succinimidyl ester (CFSE; Molecular Probes) at 37°C for 10 min. Loading was stopped by the addition of 5 % FBS, and cells were washed with PBS. CFSE-loaded cells were inoculated on a 24-well plate coated with anti-CD3/CD28 antibodies. Three days later, proliferation was measured as the dilution of CFSE dye on a FACSCalibur (BD Bioscience).

### Statistical analysis

Statistical analysis was performed using GraphPad Prism 8 software. Two-way ANOVA with Tukeýs multiple comparisons test was used for RT-qPCR data, actin polymerization, and migration assay. One-way ANOVA with Tukeýs multiple comparisons test was used for time-lapse analysis and cell homing. The number of repetitions is indicated for each experiment in the figure legend. Graphs are depicted as mean ± SEM.

## RESULTS

### RASAL3 is upregulated upon T-cell activation, localizes at the plasma membrane and binds ZAP70 and FYN

As the function of RASAL3 protein in human T cells is largely unexplored, we aimed to characterize its biology in both primary human T cells and the widely used prototypic T-cell line Jurkat (Carrasco-Padilla et al. 2023). First, we investigated how RASAL3 expression changes during T-cell activation. Therefore, we stimulated human primary T cells with a plethora of stimuli and analyzed the RASAL3 expression levels. Interestingly, human peripheral blood T cells express relatively low levels of RASAL3 at the basal state, however, its expression dramatically increases in response to the T-cell receptor (TCR) stimulation with CD3-triggering antibodies or various mitogens like phytohemagglutinin A (PHA) or the combination of phorbol ester PMA with Ionomycin. On the other hand, co-stimulatory antibodies such as anti-CD4, anti-CD28, or ICAM do not further enhance RASAL3 expression (Fig. 1A).

**Figure 1.**
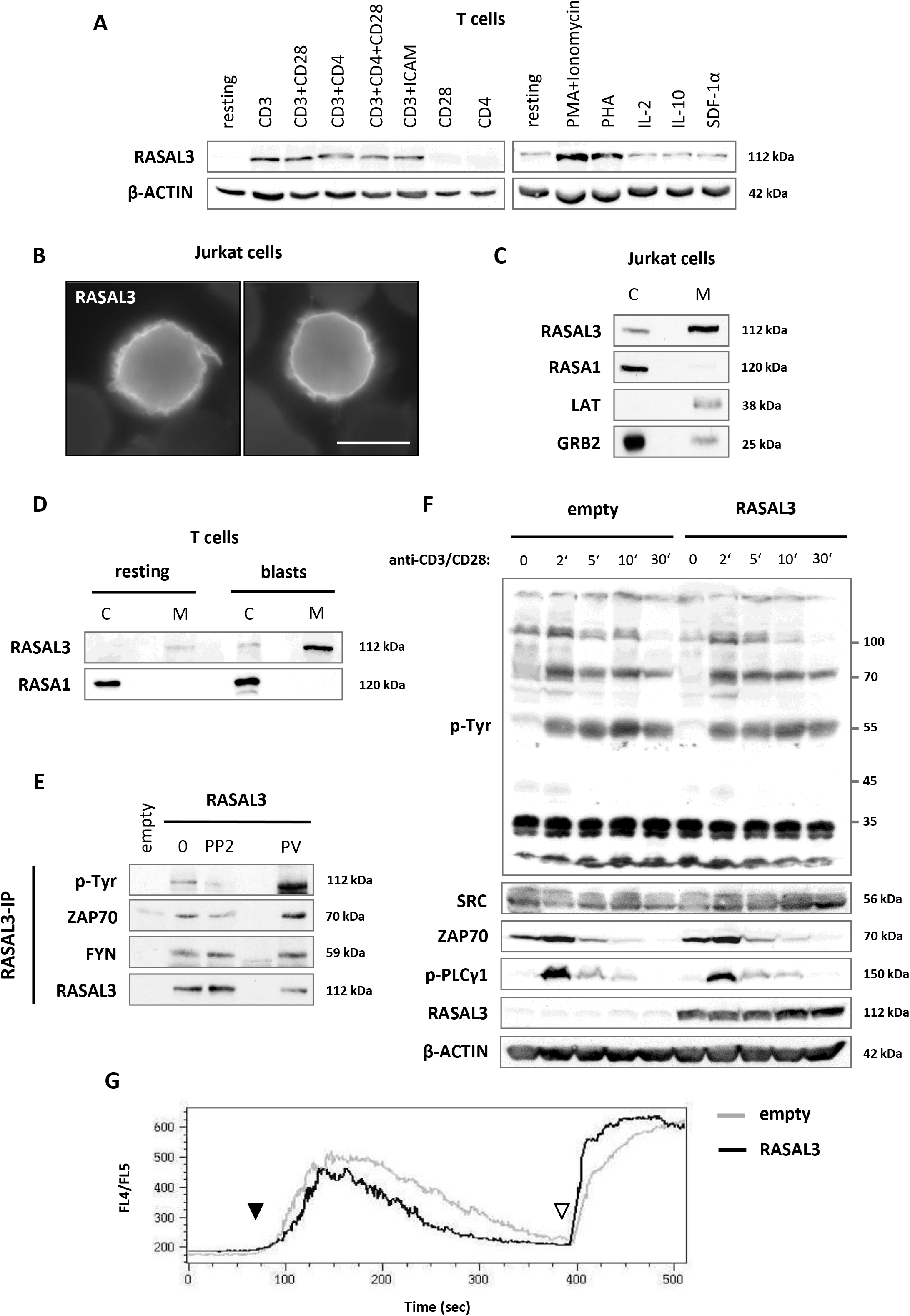
RASAL3 is upregulated upon T-cell activation, localizes at the plasma membrane, and binds ZAP70 and FYN. (A) Primary human T cells isolated from healthy donors were stimulated on plastic plates pre-coated with specified antibodies (donor 1, left panel) or by adding various soluble factors (donor 2, right panel). After 72 hours, cells were lysed and RASAL3 expression analyzed by Western blotting. β-ACTIN was used as a loading control. The experiment was repeated with multiple donors. (B) Jurkat T-cell line was transfected with RASAL3-encoding construct, and immunofluorescence staining was done to visualize RASAL3 in the cells. Scale bar = 10 µm. (C) Jurkat cells transfected with RASAL3 were fractionated into cytosolic (C) and membrane (M) parts. Western blotting shows the localization of RASAL3, RASA1 (another important RAS-GAP in T cells), LAT (Linker for Activation of T cells; membrane adaptor protein), and GRB2 (Growth Factor Receptor-Bound protein 2; mainly cytosolic adaptor protein). (D) Primary human T cells were either unstimulated (resting) or activated for 72 hours on anti-CD3/CD28-coated plastic plates (blasts). Samples were further fractionated into cytosolic (C) and membrane (M) parts, and the expression of RASAL3 and RASA1 was analyzed by Western blotting. (E) Jurkat cells were transfected with empty vector or RASAL3 plasmid. RASAL3-overexpressing cells were treated with the SFK inhibitor PP2 to prevent tyrosine phosphorylation or with pervanadate (PV) to prevent dephosphorylation. RASAL3 was immunoprecipitated and subjected to Western blotting with phospho-tyrosine antibody 4G10 (pTyr), ZAP70, FYN, and RASAL3 antibodies. (F) RASAL3-transfected or empty vector-transfected Jurkat cells were stimulated with anti-CD3/CD28 antibodies for the indicated time-points, and Western blotting was performed to assess the activation of overall protein tyrosine phosphorylation (pTyr), activation of SRC-family kinases (pY418), ZAP70, and PLC-γ. RASAL3 antibody was used to show the expression efficiency, and β-ACTIN was used as the loading control. (G) Calcium flux was measured by flow cytometry in empty vector or RASAL3-transfected Jurkat cells. Calcium flow was activated by the addition of anti-CD3 antibody (filled triangle), and maximum calcium flux was triggered by adding the calcium ionophore Ionomycin (empty triangle).

Since the RASAL3 contains PH and C2 domains (Muro et al. 2015a; Saito et al. 2015a), we determined its cellular localization in the exogenously overexpressed RASAL3 Jurkat T cells. Here, we found out that RASAL3 is primarily localized at the plasma membrane (Fig. 1B, 1C) in comparison to another important T-cell RAS-GAP RASA1, which was solely cytosolic (Fig. 1B). Membrane adaptor protein LAT (Linker for Activation of T cells) and primarily cytosolic adaptor protein GRB2 (Growth Factor Receptor-Bound Protein 2) served as markers for the membrane and cytosolic fraction, respectively (Fig. 1C). In addition, we explored the localization of endogenous RASAL3 in human primary T cells and confirmed that RASAL3 is, unlike RASA1, predominantly localized at the plasma membrane, both in resting and activated T-cells (blasts) (Fig. 1D). We then immunoprecipitated RASAL3 from Jurkat cells transfected with either empty vector or RASAL3 construct and found out that RASAL3 is tyrosine-phosphorylated, which can be prevented by treating the cells with the inhibitor of SRC-family kinases PP2 or augmented by inhibiting the protein tyrosine phosphatases via pervanadate (PV). Furthermore, RASAL3 associates with the SYK family kinase ZAP70, a tyrosine kinase critical for proper T-cell activation (Fernández-Aguilar et al. 2023), and this interaction seems to be phosphorylation-dependent. RASAL3 also binds SRC-family kinase FYN in a phosphorylation-independent manner (Fig. 1E). Since RASAL3 is localized near the plasma membrane and is associated with ZAP70, an important component of TCR signaling, we further investigated the proximal signaling induced by TCR stimulation with anti-CD3 and anti-CD28 antibodies. The overall tyrosine phosphorylation (pTyr) pattern, phosphorylation of the activatory tyrosine within the SRC-family kinases (Y418), and activation of ZAP70 or PLC gamma remained unaltered in RASAL3-overexpressing T cells compared to the control counterparts (Fig. 1F). Nevertheless, to our surprise, we observed reduced calcium flux in response to anti-CD3 triggering when RASAL3 was overexpressed (Fig. 1G).

### RASAL3 is a GTPase-activating protein for RAC/CDC42 GTPases

RASAL3 is presumed to be a GAP for small cellular GTPases, and it was originally published as a RAS-GTPase regulator (Saito et al. 2015). Nevertheless, later studies revealed its additional role in regulating the activity of the RHO-family GTPase RAC (Shin et al. 2018b). To decipher this discrepancy, we performed a series of GTPase pull-down assays to investigate the activation scope of small GTPases in response to TCR stimulation. Interestingly, we did not observe a reduction in RAS activity under RASAL3 overexpression. However, RAS activity seems to be slightly increased even though not consistently (Fig. 2A). RASAL3 overexpression also did not affect RAP1 activity (Fig. 2B). On the contrary, forced expression of RASAL3 strongly diminished the activation of both CDC42 and RAC GTPases upon anti-CD3/CD28 stimulation (Fig. 2B). Therefore, we further analyzed downstream signals resulting in the activation of mitogen-activated protein kinase (MAPK) cascades. Major MAPKs involved in T cell signaling represent ERK1/2, p38, and SAPK/JNK molecules. ERK1/2 proteins are activated by RAS-GTPase (Molina and Adjei 2006), though we did not observe any changes in phospho-ERK1/2 levels in the RASAL3-overexpressing T cells. Similarly, we did not spot any differences in p38 activation. Interestingly, anti-CD3/CD28-mediated SAPK/JNK activation, which is downstream of RAC and CDC42 GTPases (Coso et al. 1995), was strongly impaired by RASAL3 overexpression (Fig. 2C).

**Figure 2.**
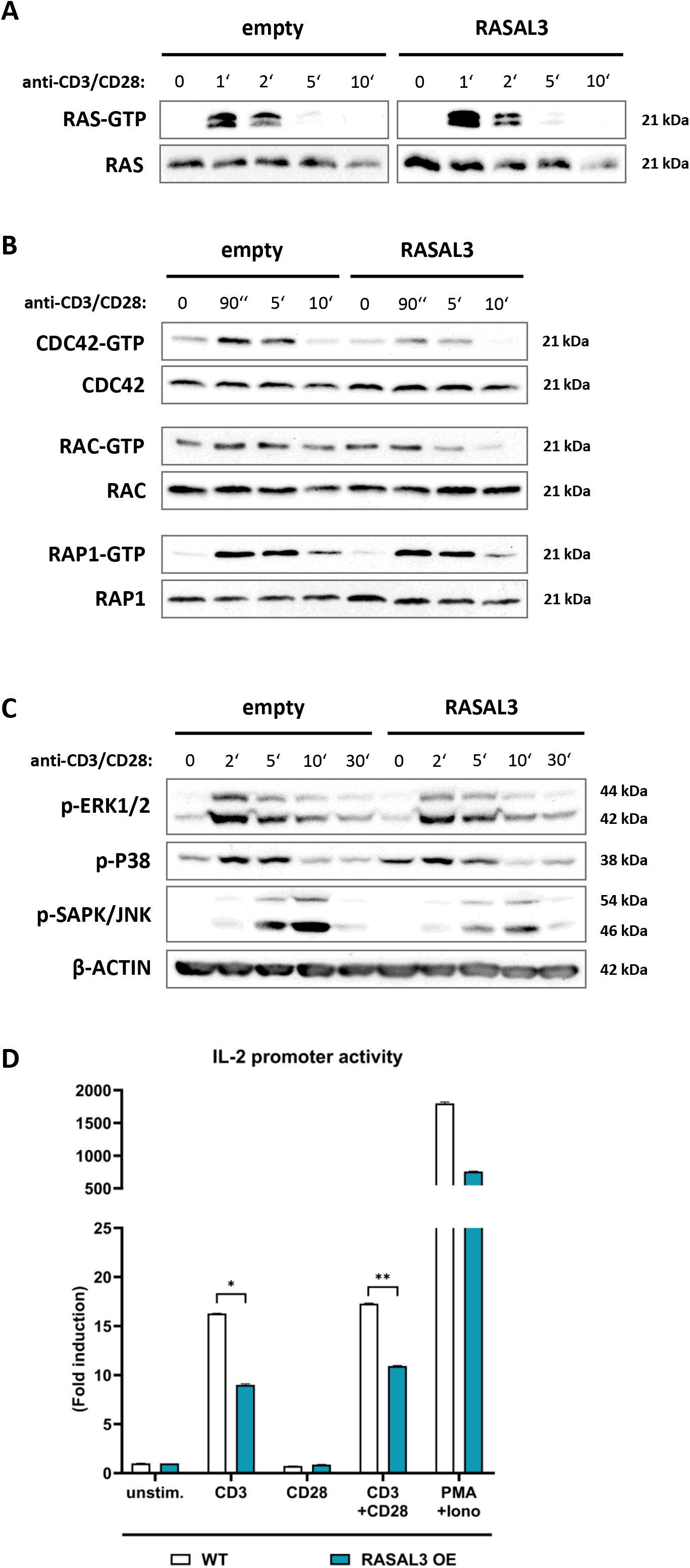
RASAL3 is a GTPase-activating protein for RAC/CDC42. (A-B) Jurkat cells transfected with empty vector or RASAL3 plasmid were stimulated with anti-CD3/CD28 antibodies for the indicated time-points and subjected to GTPase pull-down assays to assess the extent of GTP-loaded (i.e. active) form of RAS (A), CDC42, RAC, and RAP1 (B) GTPases compared to total proteins. (C) Jurkat cells transfected with empty vector or RASAL3 plasmid were stimulated with anti-CD3/CD28 antibodies for the indicated time-points and whole cell lysates were subjected to Western blotting, evaluating phosphorylation (i.e. activation) of MAP kinases ERK1/2, p38, and SAPK/JNK. β-ACTIN was used as a loading control. (D) Jurkat cells were transfected with empty vector or RASAL3 plasmid together with the Firefly luciferase reporter under the IL-2 promoter and Renilla luciferase control plasmid. Cells were stimulated with anti-CD3, anti-CD28 or anti-CD3/CD28 antibodies, with PMA+Ionomycin, or left unstimulated and subjected to dual-luciferase reporter assay. Data represent fold luminescence induction compared to unstimulated cells.

The ultimate output of T-cell receptor stimulation is the activation of the IL-2 gene promoter and IL-2 secretion (Liao et al. 2011). Therefore, we introduced an IL-2 promoter luciferase reporter construct into our RASAL3-overexpressing and control Jurkat T cells and assessed the activation of the reporter in response to T-cell stimulation. In agreement with our previous findings, RASAL3 strongly diminished activation of IL-2 promoter induced by the anti-CD3, anti-CD3/CD28, and PMA+Ionomycin stimulation (Fig. 2D).

In summary, RASAL3 functions as a GAP for RAC/CDC42 GTPases in human T cells and regulates the activation of SAPK/JNK family of MAP kinases as well as the IL-2 gene promoter activity.

### RASAL3 deficiency enhances RAC1/RAC2 and partially RAS GTPase activity, followed by increased JNK phosphorylation and elevated *c-Fos*/*c-Jun* expression

To further elucidate the importance and function of RASAL3 in T-cell signaling, we created several single-cell knockout clones. RASAL3 was knocked out in Jurkat T-cell line using CRISPR/Cas9 system (Suppl. Fig. 1A). Pull-down assays were performed in WT and RASAL3 KO Jurkat T-cell clone 1 (C1) and clone 2 (C2) following the TCR stimulation with anti-CD3/CD28 antibodies to assess the various GTPase activation kinetics. Western blot data were also quantified using ImageJ software and normalized to unstimulated WT controls. Among the analyzed GTPases, RAS exhibited the most pronounced activation in RASAL3 KO cells after short-term TCR stimulation (9-fold change/C1 and 5.9-fold change/C2 at 2 min) compared to WT cells (4.9-fold). Additionally, we observed strongly influenced RAC1 and RAC2 GTPases activity in RASAL3-deficient cells. Elevated activation of both RAC GTPases was already evident in unstimulated RASAL3 KO cells in both clones, suggesting that RASAL3 may set the activation threshold in the unstimulated cells. Moreover, RAC1 activity remained higher also during TCR stimulation and sustained RAC1 activation persisted in RASAL3 KO cells as opposed to WT cells, where a decline was observed. A comparable pattern was observed for RAC2: basal RAC2 activation was elevated in RASAL3 KO cells, and RAC2 activity was further increased and more sustained in knockout cells upon stimulation. Finally, we examined the RHO GTPase, which was only marginally affected by RASAL3 loss and inconsistent among the knockout clones (Fig. 3A).

**Figure 3.**
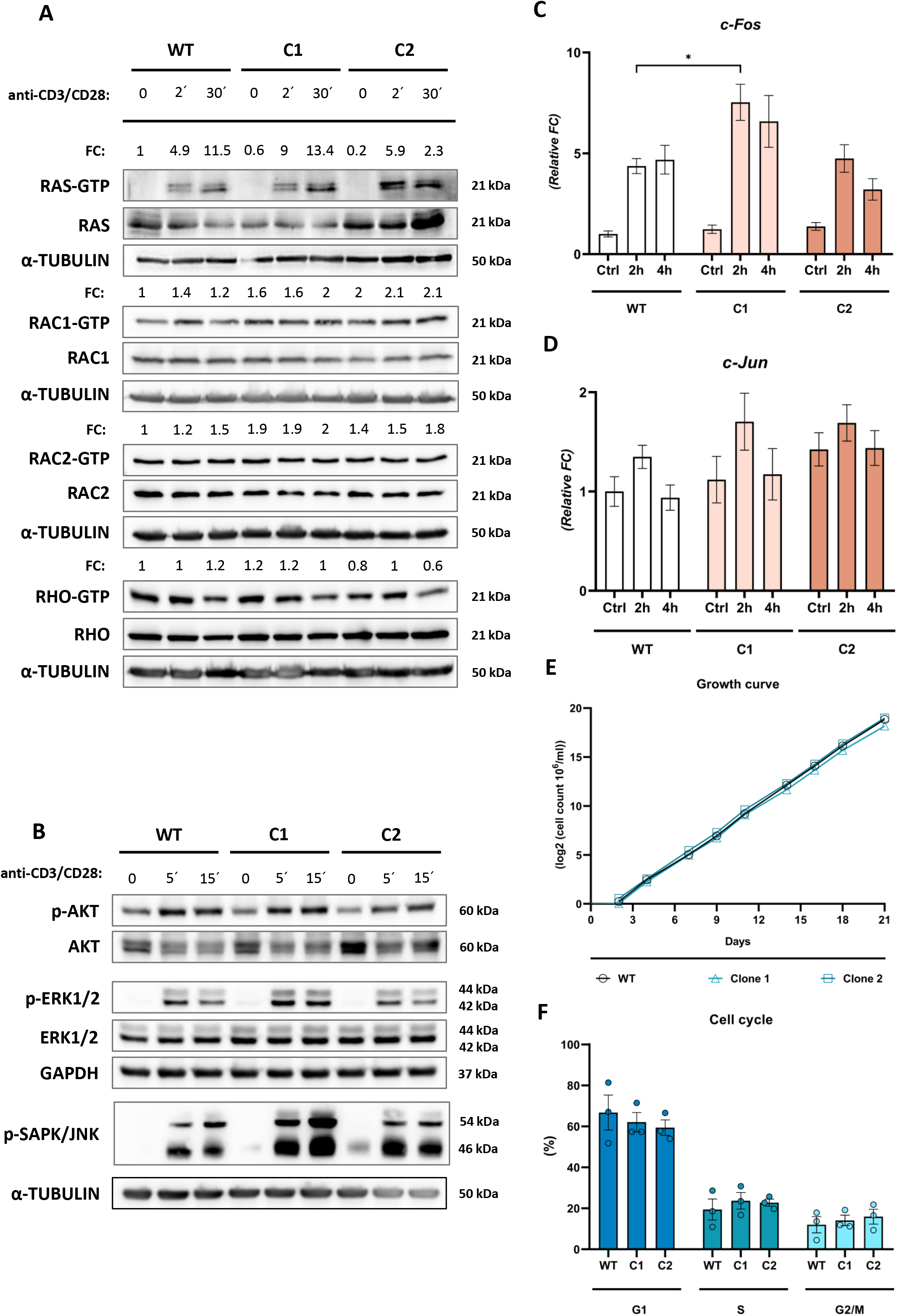
RASAL3 knockout enhances RAC1/2 GTPase activation, the SAPK/JNK phosphorylation, and Fos/Jun mRNA expression. (A) Pull-down assay for active GTPases. Wild-type (WT) and RASAL3-knockout (KO) Jurkat T cell clones 1 (C1) and 2 (C2) were stimulated with anti-CD3/CD28 antibodies for the indicated time-points. Active forms (GTP-bound) of RAS, RAC1, RAC2, and RHO were isolated using a pull-down assay kit and detected by Western blotting along with total proteins. α-TUBULIN served as a loading control. The intensity of individual bands was analyzed by densitometry using the ImageJ software, and the ratio of GTP-bound versus total GTPase form is depicted as the fold change (FC) compared to unstimulated WT cells (set as 1). (B) Jurkat WT and RASAL3 KO cells were stimulated with anti-CD3/CD28 antibodies for the indicated time-points, and levels of phosphorylated and total AKT, ERK, and SAPK/JNK proteins were analyzed by Western blotting. GAPDH and á-TUBULIN were used as loading controls. (C-D) Jurkat WT and RASAL3 KO cells were stimulated with anti-CD3/CD28 antibodies for the indicated time-points, and mRNA levels of *c-Fos* (C) and *c-Jun* (D) were measured by RT-qPCR. Data are shown as mean fold change relative to unstimulated WT (± SEM). Statistical significance is indicated as *p < 0.05. (E) WT and RASAL3 KO Jurkat cells (clones 1 and 2) were seeded in triplicate and monitored for 3 weeks. Total cell numbers were measured at each time point, and growth kinetics were plotted as the negative logarithm of mean cell counts. (F) WT and RASAL3 KO cells were collected at three different time points, stained with propidium iodide, and analyzed by flow cytometry to determine the distribution of cells in G1, S, and G2/M phases. Data are presented as mean ± SEM.

Taken together, these findings identify RASAL3 as a selective negative regulator of RAS/RAC-dependent signaling downstream of the TCR. In contrast, RHO GTPase activity remains largely unaffected, indicating that RASAL3 does not act as a global regulator of RHO family GTPases, but instead exerts specificity toward distinct signaling nodes.

To analyze the impact of RASAL3 deficiency on the downstream signaling pathways, we focused on the AKT, ERK1/2, and SAPK/JNK phosphorylation. Our data did not reveal any significant alterations in AKT phosphorylation; while ERK1/2 activation was slightly inconsistent among RASAL3 KO clones, with 1 clone increased and the other decreased compared to WT. In contrast, both RASAL3 knockout clones exhibited strongly enhanced activation of SAPK/JNK compared to control cells. Moreover, SAPK/JNK phosphorylation was already visible at the resting state, further supporting our previous notion that RASAL3 might set the activation threshold in the unstimulated cells (Fig. 3B).

Since the MAPK/ERK and SAPK/JNK signaling pathways may also regulate the expression of transcription factors *c-Fos* and *c-Jun*, we measured the mRNA levels of *c-Fos* and *c-Jun* in anti-CD3/CD28-stimulated Jurkat WT and RASAL3 KO cells using the RT-qPCR method. For both genes, maximal expression was detected at 2 hours, followed by a decline at 4 hours (Fig. 3C, 3D). *c-Fos* expression increased in RASAL3 KO clone 1 at both 2 hours (7.5-fold change, *p < 0.05) and 4 hours (6.6-fold change) compared to control cells, while KO clone 2 was more comparable to WT cells (Fig. 3C). However, the basal *c-Fos* mRNA levels were elevated in both RASAL3 KO clones (1.2-fold change in C1; 1.4-fold change in C1) than those in WT cells. The *c-Jun* expression was greater in both RASAL3-deficient clones at both time-points; however, those changes did not reach statistical significance (Fig. 3D). These data indicate that TCR stimulation also increases the expression of downstream transcription factors *c-Fos* and *c-Jun* in the absence of RASAL3.

As RASAL3 deficiency potentiated the activation of T-cell signaling pathways, we also determined whether the loss of RASAL3 affects T-cell growth and cell-cycle progression. Jurkat WT and RASAL3-KO cells (clones 1 and 2) were counted every 3–4 days over 3 weeks to generate growth curves. In parallel, cells were collected at three distinct time points and analyzed by flow cytometry to assess their distribution across the G1, S, and G2/M phases of the cell cycle. Nevertheless, no significant differences were observed in cell growth (Fig. 3E) or cell-cycle distribution (Fig. 3F). These analyses indicate that RASAL3 is dispensable for Jurkat T-cell survival and does not markedly affect their cell-cycle progression.

### RASAL3 deficiency enhances actin polymerization and promotes directed T-cell motility

Since RASAL3 was suggested to play a role in the regulation of dendritic cell migration (Olivier et al. 2024b), and we found out that RASAL3 primarily affects the activation of RAC/CDC42 GTPases, major regulators of the actin cytoskeleton (Tapon and Hall 1997), we investigated whether RASAL3 loss also affects F-actin polymerization. Thus, we quantified polymerized F-actin levels in activated Jurkat T cells stimulated with anti-CD3/CD28 antibodies. Both RASAL3 KO clones exhibited elevated basal F-actin levels compared to WT cells under unstimulated conditions (1.2-fold change in C1, Fig. 4A; 2.85-fold change in C2, Fig. 4B). The peak of the F-actin polymerization occurred after 2 minutes of stimulation in both WT and KO cells; however, this response was significantly enhanced in RASAL3 KO cells (1.8-fold change in C1, Fig. 4A; 3.6-fold change in C2, Fig. 4B) compared to WT cells. As we detected enhanced actin polymerization in RASAL3 KO T cells, we investigated actin-dependent cellular processes such as cell migration and adhesion. In the transwell migration assay, both KO clones displayed strongly increased migration towards the chemokine SDF-1α (25 % in C1 and 22 % in C2) compared to WT cells (8 %) (Fig. 4C), indicating that the loss of RASAL3 augments the migratory capacity of T-cells.

**Figure 4.**
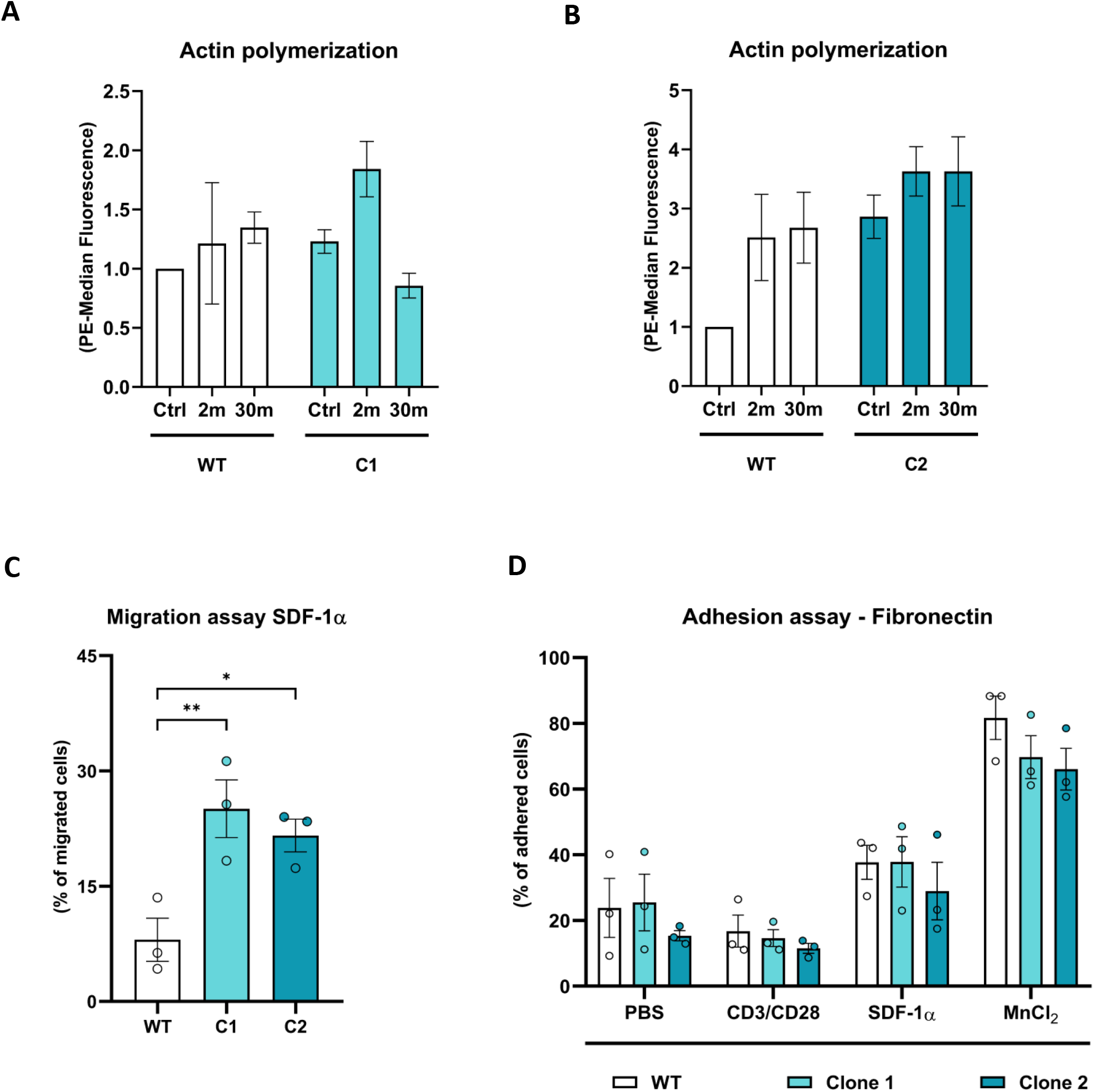
RASAL3 deficiency potentiates actin polymerization and cell migration. (A–B) Jurkat WT and RASAL3 KO cells (clones 1 and 2) were stimulated with anti-CD3/CD28 antibodies for the indicated time-points and F-actin levels were measured by flow cytometry. Data represent mean fold change relative to unstimulated WT cells (± SEM) of two independent experiments. (C) Cells were allowed to migrate toward SDF-1á for 5 hours through 81µm pore transwell inserts. Migrated cells are expressed as a percentage of the total cell input. Data represent mean ± SEM of three independent experiments with statistical significance **p < 0.01, *p < 0.05. (D) Cells were stimulated with PBS, anti-CD3/CD28 antibodies, SDF-1á, or MnCl₂ and allowed to adhere to fibronectin for 30 minutes. Adhered cells are expressed as a percentage of the total cell input. Data represent mean ± SEM of three independent experiments.

To assess the ability of RASAL3 KO Jurkat T cells to attach to the extracellular matrix components, we conducted an adhesion assay on a Fibronectin-coated surface. Cells were stimulated with MnCl2 as a non-specific stimulation (positive control), anti-CD3/CD28 antibodies for TCR activation, and SDF-1α for CXCR4 triggering. Our findings revealed an overall dropped T-cell adhesion; nevertheless, we did not observe any significant changes between WT and RASAL3 KO cells (Fig. 4D).

Based on the observed increased transwell migration in RASAL3-knockout cells, we analyzed cell migration in greater detail using time-lapse microscopy (Fig. 5A, 5B). Track displacement, defined as the distance between the first and the last cell position over time, was significantly increased in SDF-1α-stimulated RASAL3-KO cells compared to WT cells (**p < 0.01 in C1; ****p < 0.0001 in C2). Under unstimulated conditions, track displacement was also significantly elevated in clone 2 (****p < 0.0001) (Fig. 5C). The confinement ratio, reflecting directional persistence of movement (track displacement/total distance traveled), was significantly higher in SDF-1α-stimulated RASAL3-KO cells (***p < 0.001 in C1, ****p < 0.0001 in C2), as well as in unstimulated clone 2 (****p < 0.001), indicating more directed straight cell migration (Fig. 5D). Conversely, the mean directional change rate indicating more directional cell movement approached zero in both unstimulated and stimulated RASAL3-KO cells (***p < 0.001 in C2) (Fig. 5E). Consequently, the max distance traveled during the measurement period was higher in SDF-1α-stimulated RASAL3-KO cells (****p < 0.0001 in both clones) (Fig. 5F).

**Figure 5.**
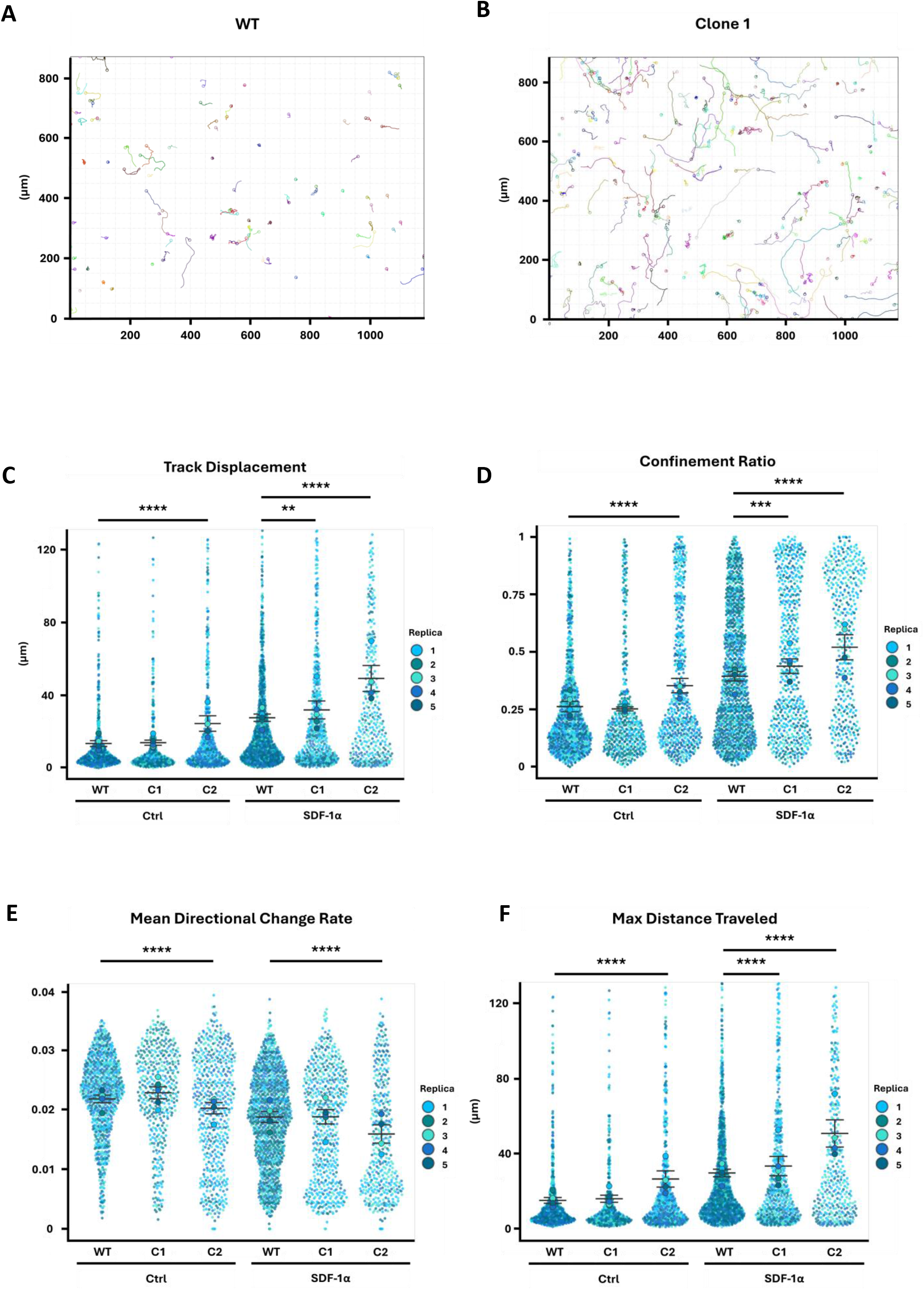
RASAL3 deficiency enhances directed cell migration. Time-lapse microscopy analysis of Jurkat cells in response to the chemokine SDF-1α. (A, B) Representative reconstructed migration trajectories of Jurkat wild-type (WT) cells and RASAL3 knockout (KO) clone 1 cells. (C–F) Quantitative parameters of cell migration: track displacement (C), confinement ratio (D), mean directional change rate (E), and maximum distance traveled (F). Data represent five independent experiments and are expressed as mean ± SEM. Statistical significance is indicated as **p < 0.01, ***p < 0.001, and ****p < 0.0001.

Taken together, consistent with the increased F-actin polymerization, the transwell migration assay and time-lapse imaging revealed significantly enhanced and more directional T cell motility of the RASAL3-deficient T cells.

### RASAL3-deficient T cells demonstrate strongly enhanced homing into the spleen *in vivo*

As RASAL3-deficient T cells displayed increased motility in vitro, we analyzed their behavior *in vivo* concerning their migration and organ-specific homing. We stained the cells with a cell tracker dye and injected a 1:1 mixture of WT and RASAL3 KO Jurkat T cells (clone 1 or 2) intraperitoneally into the immunodeficient NSG mice. We then analyzed the proportion of T cells in the blood, bone marrow, liver, and spleen by flow cytometry using anti-CD3 (FITC-conjugated) staining for T cell detection. After 48 hours, we detected the majority of T cells engrafted in the spleen, with more than 2x higher numbers of RASAL3 KO T cells compared to WT cells (Fig. 6A, 6B). To our surprise, we did not detect the cells in other organs. These data indicate that RASAL3-deficient T cells increasingly migrate *in vivo* and infiltrate the spleen faster than WT cells.

**Figure 6.**
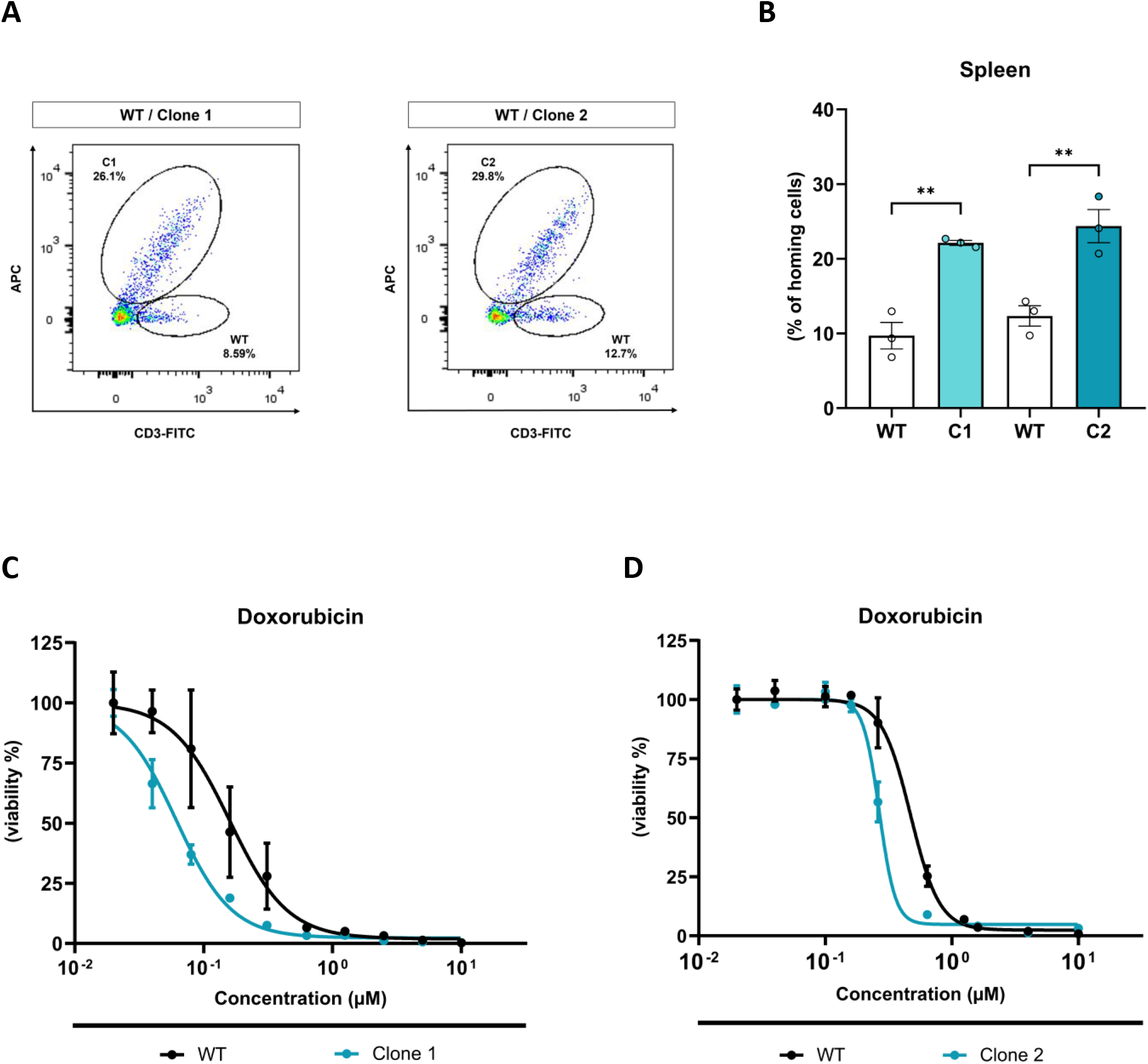
RASAL3 deficiency increases T cell homing to the spleen and sensitivity to doxorubicin. (A) Jurkat WT and RASAL3 KO clones 1 or 2 were left unstained or stained with CellTracker, alternatively, mixed 1:1 and injected intraperitoneally into three mice. Cells were isolated from the spleens 48h later, stained with anti-CD3-FITC-conjugated antibody, and the proportion of Jurkat T cell populations was analyzed by flow cytometry. Representative samples of the gating strategy are shown. (B) Cells in the spleen are expressed as a percentage of total spleen cells. Data represent mean (± SEM) from three mice. **p < 0.01. (C-D) Jurkat WT and RASAL3 KO clones 1 (C) and 2 (D) were treated with serial dilutions of doxorubicin (2-fold for clone 1; 2.5-fold for clone 2) for 72 hours. Cell viability was measured with CellTiter-Glo to generate dose–response curves. Data represent mean ± SD of three independent biological replicates.

### RASAL3-deficient Jurkat cells exhibit increased susceptibility to doxorubicin treatment

Because RASAL3 loss has been proposed as a protective factor in doxorubicin-mediated cardiotoxicity (Gao et al. 2023), we evaluated whether RASAL3 deficiency exerts a similar effect in our T-cell line. RASAL3-knockout Jurkat T cells were treated with doxorubicin for 72 h using serial dose dilutions and cell viability was assessed using the CellTiter-Glo assay. In contrast to cardiomyocytes (Gao et al. 2023), RASAL3 deficiency did not confer increased protection to doxorubicin in Jurkat T cells. Instead, RASAL3-KO cells exhibited significantly increased cell death compared to WT cells. The calculated IC₅₀ values were 0.16 µM for WT and 0.06 µM for clone 1, and 0.47 µM for WT and 0.27 µM for clone 2 (Fig. 6C, 6D), thus showing 2.6x (clone 1) and 1.7x (clone 2) higher susceptibility of RASAL3-deficient T cells to doxorubicin treatment.

### RASAL3 loss in primary human T cells unleashes the restraints on actin cytoskeleton and IL-2 gene activation

As the above experiments were primarily performed in the Jurkat T-cell line, we further assessed the consequences of RASAL3 deficiency in primary human T cells. To achieve an acute loss of RASAL3, we electroporated primary human T cells with an siRNA targeting RASAL3 (Suppl. Fig. 1A). These cells were activated on anti-CD3/CD28-antibodies-coated wells for 24 and 72 hours and the profile of their secreted cytokines was determined by a Bioplex multiplex assay (Fig. 7A and Suppl. Fig. 2). We observed that RASAL3 knockdown consistently resulted in much stronger production of IL-2 cytokine as compared to control cells (Fig. 7A). Interestingly, we did not detect any changes in any other measured cytokines such as IL-4, IL-6, IL-8, IL-10, GM-CSF, IFN-γ and TNF-α (Suppl. Fig. 2). To determine whether increased IL-2 production results also in increased T-cell proliferation, we labeled the cells with CFSE and assessed its serial dilution throughout cell divisions. Indeed, RASAL3-knockdown cells displayed faster CFSE dilution and a much higher proportion of knockdown cells proliferated three or more times as compared to control cells (Fig. 7B). As we have demonstrated the major impact of RASAL3 on actin cytoskeleton and actin-dependent migration, we also evaluated these processes in primary T cells transfected with RASAL3 siRNA. As expected, RASAL3-knockdown cells showed a higher actin polymerization rate (Fig. 7C) and increased cell migration in response to multiple chemokines, such as SDF-1α, CCL19, and CCL21 (Fig. 7D).

**Figure 7.**
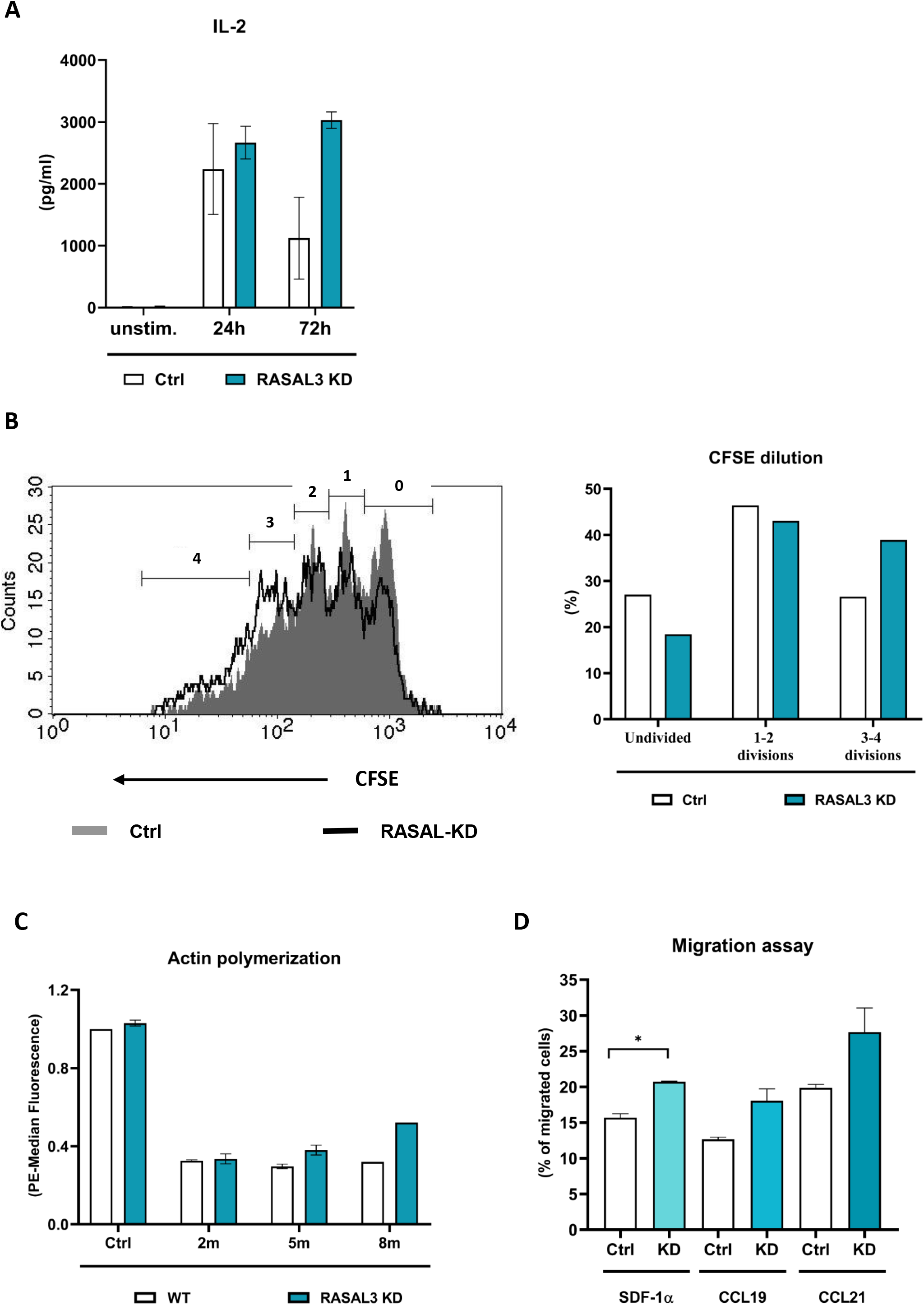
RASAL3 knockdown in primary human T cells enhances IL-2 production, cell proliferation, and cell migration. (A) Primary human T cells were transfected with RASAL3 siRNA or scrambled control and activated on anti-CD3/CD28-coated plastic plates for 24 and 72 hours. IL-2 production was determined by flow cytometry using the Bioplex 8-plex assay. Depicted is mean (± SEM) from three independent biological replicates from three different T-cell donors. (B) Primary human T cells transfected with RASAL3 siRNA or scrambled control were labeled with CFSE and cultured on anti-CD3/CD28-coated plastic plates for 72 hours. CFSE signal dilution throughout cell divisions was determined by flow cytometry. A representative example is shown. The right panel shows the percentage of cells that underwent defined number of divisions. (C) WT and RASAL3-knockdown (KD) primary human T cells were stimulated with anti-CD3/CD28 antibodies for the indicated time-points and F-actin levels were measured by flow cytometry. Data represent mean fold change relative to unstimulated WT cells (± SEM) of three independent experiments. (D) Transwell migration of control and RASAL3-knockdown (KD) primary human T cells towards the chemokines SDF-1á, CCL19, and CCL21. Migrated cells are expressed as a percentage of the total cell input. Data represent mean ± SEM of three independent experiments.

In summary, reducing RASAL3 protein levels in primary human T cells results in stronger actin polymerization and cell migration. At the same time, these cells display increased activation of IL-2 gene expression and enhanced cell proliferation.

Collectively, these findings suggest that RASAL3 functions predominantly as a GAP for RAC/CDC42 GTPases, and thereby attenuating SAPK/JNK signaling involved in IL-2 gene activation. Concurrently, RASAL3 restricts actin cytoskeleton polymerization, thereby controlling cell motility in response to chemokine gradients.

## DISCUSSION

This study provides a comprehensive analysis of the role of RASAL3 in human T-cell signaling and cellular functions using the Jurkat T-cell acute lymphoblastic leukemia (T-ALL) cell line as the commonly used T-cell model (Carrasco-Padilla et al. 2023), along with primary human T cells. Our results demonstrate that RASAL3 regulates small GTPase activity and their downstream signaling pathways leading to IL-2 gene activity, reshapes cytoskeletal dynamics, migratory behavior, and modulates cellular response to chemotherapeutic stress. Taken together, these findings position RASAL3 as an important regulator of signaling amplitude and functional plasticity in T cells.

Mechanistically, RASAL3 functions as a GTPase-activating protein (GAP) that limits both the intensity and duration of signaling mediated by the RAS-GTPase superfamily. Although RASAL3 was originally identified as a RAS-GAP in murine T cells (Saito et al. 2015; Muro et al. 2015), subsequent work by Shin et al. proposed that RASAL3 predominantly activates RAC2 GTPase, displaying only minimal activity toward RAS (Shin et al. 2018). Nevertheless, the aforementioned study focused exclusively on the inactive state of Jurkat cells, whereas our data captured the dynamics of cellular responses during T-cell activation. Our findings in RASAL3-deficient Jurkat cells confirm previous studies indicating the effect of RASAL3 on the RAS GTPase activity (Saito et al. 2015; Muro et al. 2015) and further expand them by demonstrating substantial regulatory impact upon RAC family GTPases. Interestingly, RASAL3 overexpression did not cause any reduction in RAS GTPase activation, while the activities of RAC and CDC42 GTPases were markedly affected, further supporting a prominent role for RASAL3 in the regulation of RAC-family signaling. In agreement with previous findings, neither RAP1 nor RHO GTPases appear to be regulated by RASAL3 in T lymphocytes, in contrast to dendritic cells (Olivier et al. 2024). Together, these findings suggest that RASAL3 primarily modulates signaling downstream of proximal TCR activation by selectively regulating members of the RAS-GTPase superfamily.

Consistent with this notion, we did not observe any alterations in proximal TCR signaling events, including global tyrosine phosphorylation, ZAP70 activation, or PLC-γ phosphorylation. However, RASAL3 overexpression attenuated calcium flux induced by anti-CD3 stimulation, indicating that RASAL3 may influence calcium signaling independently of proximal TCR signaling pathways, although the underlying mechanism remains unclear and requires further investigation.

One possible explanation may be related to the domain organization and subcellular localization of RASAL3. The protein contains a C2 interaction domain (Saito et al. 2015), which is thought to mediate calcium-dependent targeting of proteins to the plasma membrane. Therefore, fluctuations in intracellular calcium levels may not only represent a downstream consequence of T-cell activation but could also regulate the subcellular localization and activity of RASAL3 itself (Corbin et al. 2007). Accordingly, we found that RASAL3 is predominantly localized at the plasma membrane, where it is presumably recruited through its PH and C2 domains. At the plasma membrane, RASAL3 may interact with FYN, a Src-family kinase associated with the T-cell receptor complex (Takeuchi et al. 1993), and consequently undergo tyrosine phosphorylation. Such membrane localization would place RASAL3 in close proximity to signaling complexes generated upon TCR engagement and provide a mechanistic basis for its ability to modulate the activity of RAS and RAC family GTPases.

In agreement with this model, altered RASAL3 expression in TCR-stimulated T cells modulated the activation of SAPK/JNK, a downstream effector of RAC/CDC42 signaling (Coso et al. 1995), and, to a lesser extent, ERK1/2, a key component of the RAS signaling pathway (Chang et al. 2003). In contrast, no significant changes were observed in the activation of AKT or p38 MAPK. These findings are consistent with our GTPase activity data and suggest that RASAL3 selectively regulates signaling pathways downstream of RAS and RAC family GTPases.

Upon TCR stimulation through CD3 and CD28 co-stimulation, the RAS–ERK and RAC/CDC42–JNK signaling cascades are activated and converge on the formation of the AP-1 transcription factor complex, ultimately leading to IL-2 gene transcription (Faris et al. 1996). AP-1 is primarily composed of dimers formed by members of the Fos, Jun, Maf, and ATF protein families and regulates a broad spectrum of cellular processes, including proliferation, differentiation, and apoptosis (Hess et al. 2004). In T lymphocytes, AP-1 plays a central role in IL-2 promoter activation and T-cell lineage differentiation (Jain et al. 1992). Therefore, the observed effects of RASAL3 on ERK1/2 and JNK signaling suggested that RASAL3 may influence T-cell activation through modulation of AP-1-dependent transcriptional programs.

To test this possibility, we examined the expression of the AP-1 components *c-Fos* and *c-Jun*, whose expression is regulated by MAPK signaling pathways (Monje et al. 2005; Johnson and Nakamura 2007). Following anti-CD3/CD28 stimulation, RASAL3-deficient Jurkat cells exhibited substantially increased expression of both *c-Fos* and *c-Jun*, supporting possible enhanced AP-1 activity. Consistent with these observations, RASAL3 overexpression reduced IL-2 promoter activity, whereas RASAL3 knockdown markedly increased IL-2 production in primary T lymphocytes. Together, these results identify RASAL3 as a negative regulator of the signaling network linking RAS/RAC activation to AP-1-mediated IL-2 transcription.

Since IL-2 is a key autocrine growth factor for activated T cells (Toribio et al. 1989), we next examined whether altered RASAL3 expression also affects proliferative responses. Indeed, RASAL3 knockdown enhanced the proliferation of primary T cells, most likely as a consequence of increased IL-2 production. Notably, we were unable to detect measurable IL-2 production in the Jurkat T-cell line used in our study (data not shown). Consistent with this observation, no significant differences in proliferation or cell-cycle distribution were detected in RASAL3-deficient Jurkat cells, which are immortalized and constitutively proliferating. Interestingly, these findings differ from previous studies in mice, where RASAL3 deficiency impaired proliferation and promoted apoptosis of naïve T cells and natural killer T (NK-T) cells through reduced expression of the anti-apoptotic protein BCL-2 (Muro et al. 2015; Shin et al. 2018). Taken together, these results suggest that the role of RASAL3 in regulating proliferation and survival is context- and stage-dependent and may differ between species and/or be determined by microenvironmental interactions *in vivo*.

Apart from the transcriptional signaling alterations, RASAL3 significantly affected cytoskeletal organization and migratory behavior. RAC1, RAC2, and CDC42 are key regulators of actin polymerization and directional migration in T cells (Raftopoulou and Hall 2004). In agreement with enhanced RAC1/2 and CDC42 activation, RASAL3-deficient Jurkat cells exhibited increased actin polymerization and enhanced migration toward stromal cell–derived factor 1 alpha (SDF-1α). SDF-1α is an important T-cell chemoattractant involved in numerous biological processes, including hematopoiesis, angiogenesis, organogenesis, osteogenesis, tissue repair, and tumor metastasis. SDF-1α primarily signals through the C-X-C chemokine receptor type 4 (CXCR4) and modulates immune responses by recruiting immune cells to sites of inflammation, infection, and tumor (Dunussi-Joannopoulos et al. 2002; Biyani et al. 2024). The enhanced migratory response observed in RASAL3-deficient cells suggests that loss of RASAL3 lowers the activation threshold for chemokine-induced RAC/CDC42 signaling and cytoskeletal rearrangements. Importantly, the absence of RASAL3 was associated with more directed cell movement and fewer directional changes, suggesting a potential role in regulating cell polarity and actin filament branching.

Interestingly, despite the central role of TCR and integrin signaling in adhesion and immunological synapse formation (O’Rourke and Mescher 1990; Dustin and Springer 1989; Bertoni et al. 2018), RASAL3 deficiency did not significantly affect Jurkat T-cell adhesion to fibronectin (FN). FN is a high-molecular-weight extracellular matrix glycoprotein that binds to integrins, particularly very late antigen-4 (VLA-4; integrin α4β1) and lymphocyte function-associated antigen-1 (LFA-1; integrin αLβ2), and plays a crucial role in T-cell adhesion, migration, and activation (Rodriguez et al. 1992; Dalton and Lemmon 2021). The absence of significant changes in FN-mediated adhesion suggests that RASAL3 preferentially regulates migratory rather than adhesive functions of T cells, potentially by selectively modulating RAC/CDC42-dependent cytoskeletal dynamics without substantially altering integrin-mediated adhesion.

The functional relevance of this phenotype is further supported by the increased homing of RASAL3-deficient T cells to the spleen. The spleen is a major secondary lymphoid organ responsible for immune surveillance of blood-borne antigens, T- and B-cell activation, and orchestration of adaptive immune responses (Lewis et al. 2019; Bronte and Pittet 2013). The splenic microenvironment is characterized by high expression of chemokines, adhesion molecules, and stromal signals that regulate lymphocyte trafficking and retention (Mueller and Germain 2009). Enhanced accumulation of RASAL3-deficient T cells in the spleen may therefore reflect increased responsiveness to chemokine gradients, particularly the SDF-1α/CXCR4 axis, as well as enhanced migratory capacity and amplified signaling downstream of RAC and CDC42 pathways, promoting more efficient interaction with the splenic microenvironment. Importantly, enhanced activation of GTPases accompanied by augmented motility of RASAL3-deficient cells was also described for dendritic cells (Olivier et al., 2024).

Finally, we investigated the impact of RASAL3 deficiency on cellular responses to doxorubicin, an anthracycline chemotherapeutic agent widely used in the treatment of hematological malignancies, including acute lymphoblastic leukemia. Doxorubicin exerts cytotoxic effects through DNA intercalation, inhibition of topoisomerase II, induction of DNA double-strand breaks, and generation of reactive oxygen species (ROS), ultimately leading to apoptosis (Johnson-Arbor et al. 2026). While reduced RASAL3 expression has been reported to protect cardiomyocytes from doxorubicin-induced toxicity (Gao et al. 2023), we observed the opposite effect in Jurkat T cells, where RASAL3 deficiency increased sensitivity to doxorubicin-induced cell death. One possible explanation is that enhanced signaling in RASAL3-deficient T cells promotes a more metabolically active and proliferative state, rendering these cells more vulnerable to genotoxic stress. Altered JNK signaling may further shift the balance between survival and apoptosis in response to DNA damage (J. Liu and Lin 2005). Moreover, RASAL3 mutations have been linked to synovial sarcoma (Qi et al. 2015), and its epigenetic silencing has been proposed as a sensor of metabolic reprogramming in prostate cancer (Mishra et al., n.d.), highlighting RASAL3 as a potential therapeutic target for enhancing chemotherapy efficacy. However, these findings require validation in future studies.

The signaling and functional alterations observed in RASAL3-deficient T cells raise the question of whether modulating RASAL3 could have implications for cellular immunotherapies. Enhanced T-cell functionality is particularly relevant in the context of CAR-T cell therapy, where improving the signaling capacity, persistence, and migratory potential of engineered T cells remains a major challenge. Chimeric antigen receptor (CAR) T cells express genetically modified receptors combining an antigen-binding domain derived from monoclonal antibodies with the TCR domains, enabling recognition of tumor-associated antigens, mediating CAR-T cell activation and tumor cell eradication (Zugasti et al. 2025). Insufficient CAR-T cell persistence and the occurrence of severe adverse effects, including cytokine release syndrome, immune effector cell–associated neurotoxicity syndrome, and immune effector cell-associated haematotoxicity (Adkins 2019), highlight the need for a deeper understanding of molecular mechanisms that regulate T-cell function.

Enhanced TCR-triggered signaling, along with increased IL-2 secretion and motility in RASAL3-deficient T cells, could theoretically improve CAR-T cell activation and migration toward tumor-derived chemokines, such as SDF-1α. However, excessive activation of these pathways may also promote premature exhaustion, altered differentiation states, or increased susceptibility to stress-induced apoptosis. Therefore, RASAL3 deficiency may represent a double-edged regulatory mechanism, potentially enhancing effector and migratory functions while perhaps compromising long-term stability and persistence of engineered T cells.

In conclusion, our data identify RASAL3 as a significant modulator of small GTPase signaling and downstream functional responses in human T cells. Rather than acting as a universal regulator of proliferation or survival, RASAL3 fine-tunes signaling intensity, cytoskeletal dynamics, migratory behavior, and stress sensitivity in a context-dependent manner. Interestingly, although RASAL3 expression is relatively low in resting primary human T cells, it becomes markedly upregulated upon T-cell activation. We therefore speculate that the primary role of RASAL3 may be to modulate the magnitude of T-cell activation and contribute to the termination of T-cell–mediated responses. Our findings provide new insights into the role of RASAL3 in human T-cell biology and highlight its potential relevance for understanding signaling dysregulation in T-cell malignancies and for the development of immunotherapeutic strategies.

## Declaration of generative AI and AI-assisted technologies in the manuscript preparation process

During the preparation of this work the author(s) used ChatGPT (OpenAI, San Francisco, USA) and Grammarly (Grammarly Inc., San Francisco, USA) for language editing assistance and stylistic improvements of the manuscript. After using this tool/service, the author(s) reviewed and edited the content as needed and take(s) full responsibility for the content of the published article.

## AUTHOR CONTRIBUTIONS

**S. Varadínková:** Conceptualization, Methodology, Investigation, Data curation, Writing of the manuscript, Visualization, Project administration; **P. Oslacký, K. Kvasničková, P. Cigánková, N. V. Gottumukkala:** Investigation, Š. **Čada:** Investigation, Data curation; **J. A. Lindquist:** Conceptualization, Resources, Supervision; **M. Šmída:** Conceptualization, Methodology, Investigation, Resources, Writing of the manuscript, Supervision, Project administration, Funding acquisition

## ACKNOWLEDGEMENTS (all funding sources)

We acknowledge the Core Facility Genomics (CEITEC) supported by the NCMG research infrastructure (LM2023067 funded by MEYS CR) and Flow cytometry laboratory at CEITEC MU supported by the EATRIS-CZ research infrastructure (LM2023053 funded by MEYS CR) for their support with obtaining scientific data presented in this paper. This study was funded by the research grant from the Czech Science Foundation (project no. 22-35273S), the project National Institute for Cancer Research (Programme EXCELES, ID Project No. LX22NPO5102) - Funded by the European Union – Next Generation EU, and project MUNI/A/1733/2025.

---

**Supplementary Figure 1. RASAL3 knockout in Jurkat cells and knockdown in primary T cells**

(A) RASAL3 protein expression in Jurkat wild-type (WT) cells and RASAL3 knockout (KO) clones (C1 and C2) was determined by Western blot analysis (left panel). The right panel shows RASAL3 protein knockdown efficiency in primary human T cells. á-TUBULIN and â-ACTIN served as loading controls.

(B) Sequences of primers used for RT-qPCR analysis. All primers were synthesized and purchased from Generi Biotech.

(C) Sequences of single-guide RNAs (sgRNAs) used for electroporation to generate RASAL3 KO Jurkat T cells.

**Supplementary Figure 2. Cytokine production in primary human T cells following RASAL3 knockdown**

Primary human T cells were transfected with RASAL3 siRNA or scrambled control siRNA and activated on anti-CD3/CD28-precoated plates for 24 or 72 h. Production of IL-2 (A), IL-4 (B), IL-6 (C), IL-8 (D), IL-10 (E), GM-CSF (F), IFN-γ (G), and TNF-α (H) was quantified by flow cytometry using the Bio-Plex 8-plex assay. Data are presented as mean ± SEM from three independent biological replicates derived from three different T-cell donors.

**Supplementary Figure 3. Antibodies used for Western blot analysis**

List of antibodies used for Western blot analysis, including molecular weight, dilution, host species, and manufacturer. Manufacturers: Aviva Systems Biology (ASB), Cell Signaling Technology (CST), Proteintech (P), Sigma-Aldrich (SA), and Thermo Fisher Scientific (TFS).

## REFERENCES

Adkins, Sherry. 2019. ‘CAR T-Cell Therapy: Adverse Events and Management’. Journal of the Advanced Practitioner in Oncology 10 (Suppl 3): 21–28. 10.6004/jadpro.2019.10.4.11.

Bertoni, Alessandra, Oscar Alabiso, Alessandra Silvia Galetto, and Gianluca Baldanzi. 2018. ‘Integrins in T Cell Physiology’. International Journal of Molecular Sciences 19 (2): 485. 10.3390/ijms19020485.

Biyani, Shruti, Amol Patil, and Vinit Swami. 2024. ‘The Influence of SDF-1 (CXCL12) Gene in Health and Disease: A Review of Literature’. Biophysical Reviews 17 (1): 127–38. 10.1007/s12551-024-01230-5.

Bronte, Vincenzo, and Mikael J. Pittet. 2013. ‘The Spleen in Local and Systemic Regulation of Immunity’. Immunity 39 (5): 806–18. 10.1016/j.immuni.2013.10.010.

Čada, Štěpán, Olga Vondálová Blanářová, Kristína Gömoryová, et al. 2022. ‘Role of Casein Kinase 1 in the Amoeboid Migration of B-Cell Leukemic and Lymphoma Cells: A Quantitative Live Imaging in the Confined Environment’. Frontiers in Cell and Developmental Biology 10 (December). 10.3389/fcell.2022.911966.

Carrasco-Padilla, C., O. Aguilar-Sopeña, Alvaro Gómez-Morón, et al. 2023. ‘T Cell Activation and Effector Function in the Human Jurkat T Cell Model’. Methods in Cell Biology 178: 25–41. 10.1016/bs.mcb.2022.09.012.

Chang, F., L. S. Steelman, J. T. Lee, et al. 2003. ‘Signal Transduction Mediated by the Ras/Raf/MEK/ERK Pathway from Cytokine Receptors to Transcription Factors: Potential Targeting for Therapeutic Intervention’. Leukemia 17 (7): 1263–93. 10.1038/sj.leu.2402945.

Cherfils, Jacqueline, and Mahel Zeghouf. 2013. ‘Regulation of Small GTPases by GEFs, GAPs, and GDIs’. Physiological Reviews 93 (1): 269–309. 10.1152/physrev.00003.2012.

Corbin, John A., John H. Evans, Kyle E. Landgraf, and Joseph J. Falke. 2007. ‘Mechanism of Specific Membrane Targeting by C2 Domains: Localized Pools of Target Lipids Enhance Ca2+ Affinity’. Biochemistry 46 (14): 4322–36. 10.1021/bi062140c.

Coso, O. A., M. Chiariello, J. C. Yu, et al. 1995. ‘The Small GTP-Binding Proteins Rac1 and Cdc42 Regulate the Activity of the JNK/SAPK Signaling Pathway’. Cell 81 (7): 1137–46. 10.1016/s0092-8674(05)80018-2.

Dalton, Caleb J., and Christopher A. Lemmon. 2021. ‘Fibronectin: Molecular Structure, Fibrillar Structure and Mechanochemical Signaling’. Cells 10 (9): 2443. 10.3390/cells10092443.

Dunussi-Joannopoulos, Kyriaki, Krystyna Zuberek, Kathlene Runyon, et al. 2002. ‘Efficacious Immunomodulatory Activity of the Chemokine Stromal Cell-Derived Factor 1 (SDF-1): Local Secretion of SDF-1 at the Tumor Site Serves as T-Cell Chemoattractant and Mediates T-Cell-Dependent Antitumor Responses’. Blood 100 (5): 1551–58.

Dustin, Michael L., and Timothy A. Springer. 1989. ‘T-Cell Receptor Cross-Linking Transiently Stimulates Adhesiveness through LFA-1’. Nature 341 (6243): 619–24. 10.1038/341619a0.

Faris, M., N. Kokot, L. Lee, and A. E. Nel. 1996. ‘Regulation of Interleukin-2 Transcription by Inducible Stable Expression of Dominant Negative and Dominant Active Mitogen-Activated Protein Kinase Kinase Kinase in Jurkat T Cells. Evidence for the Importance of Ras in a Pathway That Is Controlled by Dual Receptor Stimulation’. The Journal of Biological Chemistry 271 (44): 27366–73. 10.1074/jbc.271.44.27366.

Fernández-Aguilar, Luis M., Inmaculada Vico-Barranco, Mikel M. Arbulo-Echevarria, and Enrique Aguado. 2023. ‘A Story of Kinases and Adaptors: The Role of Lck, ZAP-70 and LAT in Switch Panel Governing T-Cell Development and Activation’. Biology 12 (9): 1163. 10.3390/biology12091163.

Gao, Ri-Feng, Kun Yang, Ya-Nan Qu, et al. 2023a. ‘m6A Demethylase ALKBH5 Attenuates Doxorubicin-Induced Cardiotoxicity via Posttranscriptional Stabilization of Rasal3’. iScience 26 (3): 106215. 10.1016/j.isci.2023.106215.

Gao, Ri-Feng, Kun Yang, Ya-Nan Qu, et al. 2023b. ‘m6A Demethylase ALKBH5 Attenuates Doxorubicin-Induced Cardiotoxicity via Posttranscriptional Stabilization of Rasal3’. iScience 26 (3): 106215. 10.1016/j.isci.2023.106215.

Hess, Jochen, Peter Angel, and Marina Schorpp-Kistner. 2004. ‘AP-1 Subunits: Quarrel and Harmony among Siblings’. Journal of Cell Science 117 (25): 5965–73. 10.1242/jcs.01589.

Hong, Changwan, Megan A. Luckey, and Jung-Hyun Park. 2012. ‘Intrathymic IL-7: The Where, When, and Why of IL-7 Signaling during T Cell Development’. Seminars in Immunology, IL-7 in the Immune System: Development and Function, vol. 24 (3): 151–58. 10.1016/j.smim.2012.02.002.

Jain, J., V. E. Valge-Archer, and A. Rao. 1992. ‘Analysis of the AP-1 Sites in the IL-2 Promoter’. Journal of Immunology (Baltimore, Md.1: 1950) 148 (4): 1240–50.

Johnson, Gary L., and Kazuhiro Nakamura. 2007. ‘The C-Jun Kinase/Stress-Activated Pathway: Regulation, Function and Role in Human Disease’. *Biochimica et Biophysica Acta (BBA) - Molecular Cell Research*, Mitogen-Activated Protein Kinases: New Insights on Regulation, Function and Role in Human Disease, vol. 1773 (8): 1341–48. 10.1016/j.bbamcr.2006.12.009.

Johnson-Arbor, Kelly, Steven Douedi, and Preeti Patel. 2026. ‘Doxorubicin’. In StatPearls. StatPearls Publishing. http://www.ncbi.nlm.nih.gov/books/NBK459232/.

Lewis, Steven M., Adam Williams, and Stephanie C. Eisenbarth. 2019. ‘Structure-Function of the Immune System in the Spleen’. Science Immunology 4 (33): eaau6085. 10.1126/sciimmunol.aau6085.

Liang, Mei, Xiangzhi Meng, Boxuan Zhou, and Yushun Gao. 2022. ‘RASAL3 Predicts Overall Survival and CD8+ T Lymphocyte Infiltration in Lung Adenocarcinoma’. Journal of Cellular and Molecular Medicine 26 (24): 6056–65. 10.1111/jcmm.17625.

Liao, Wei, Jian-Xin Lin, and Warren J. Leonard. 2011. ‘IL-2 Family Cytokines: New Insights into the Complex Roles of IL-2 as a Broad Regulator of T Helper Cell Differentiation’. Current Opinion in Immunology 23 (5): 598–604. 10.1016/j.coi.2011.08.003.

Lin, Yiyan, Dhiman Sankar Pal, Parijat Banerjee, et al. 2024. ‘Ras Suppression Potentiates Rear Actomyosin Contractility-Driven Cell Polarization and Migration’. Nature Cell Biology 26 (7): 1062–76. 10.1038/s41556-024-01453-4.

Lin, Zigen, Xiaozhu Tang, Yuhao Cao, et al. 2022. ‘CD229 Interacts with RASAL3 to Activate RAS/ERK Pathway in Multiple Myeloma Proliferation’. Aging 14 (22): 9264–79. 10.18632/aging.204405.

Liu, Jing, and Anning Lin. 2005. ‘Role of JNK Activation in Apoptosis: A Double-Edged Sword’. Cell Research 15 (1): 36–42. 10.1038/sj.cr.7290262.

Liu, Zhaoyun, Wenhui Lei, Hao Wang, Xiaohan Liu, and Rong Fu. 2024. ‘Challenges and Strategies Associated with CAR-T Cell Therapy in Blood Malignancies’. Experimental Hematology & Oncology 13 (February): 22. 10.1186/s40164-024-00490-x.

Mishra, Rajeev, Subhash Haldar, Veronica Placencio, et al. n.d. ‘Stromal Epigenetic Alterations Drive Metabolic and Neuroendocrine Prostate Cancer Reprogramming’. The Journal of Clinical Investigation 128 (10): 4472–84. 10.1172/JCI99397.

Molina, Julian R., and Alex A. Adjei. 2006. ‘The Ras/Raf/MAPK Pathway’. Journal of Thoracic Oncology 1 (1): 7–9. 10.1016/S1556-0864(15)31506-9.

Monje, Paula, Javier Hernández-Losa, Ruth J. Lyons, Maria D. Castellone, and J. Silvio Gutkind. 2005. ‘Regulation of the Transcriptional Activity of C-Fos by ERK. A Novel Role for the Prolyl Isomerase PIN1’. The Journal of Biological Chemistry 280 (42): 35081–84. 10.1074/jbc.C500353200.

Mueller, Scott N., and Ronald N. Germain. 2009. ‘Stromal Cell Contributions to the Homeostasis and Functionality of the Immune System’. Nature Reviews. Immunology 9 (9): 618–29. 10.1038/nri2588.

Muro, Ryunosuke, Takeshi Nitta, Masayuki Kitajima, Toshiyuki Okada, and Harumi Suzuki. 2018. ‘Rasal3-Mediated T Cell Survival Is Essential for Inflammatory Responses’. Biochemical and Biophysical Research Communications 496 (1): 25–30.

Muro, Ryunosuke, Takeshi Nitta, Toshiyuki Okada, Hitoshi Ideta, Takeshi Tsubata, and Harumi Suzuki. 2015a. ‘The Ras GTPase-Activating Protein Rasal3 Supports Survival of Naive T Cells’. PloS One 10 (3): e0119898. 10.1371/journal.pone.0119898.

Muro, Ryunosuke, Takeshi Nitta, Toshiyuki Okada, Hitoshi Ideta, Takeshi Tsubata, and Harumi Suzuki. 2015b. ‘The Ras GTPase-Activating Protein Rasal3 Supports Survival of Naive T Cells’. PloS One 10 (3): e0119898. 10.1371/journal.pone.0119898.

Olivier, Jean-Frederic, David Langlais, Thiviya Jeyakumar, et al. 2024a. ‘CCDC88B Interacts with RASAL3 and ARHGEF2 and Regulates Dendritic Cell Function in Neuroinflammation and Colitis’. Communications Biology 7 (1): 77. 10.1038/s42003-023-05751-9.

Olivier, Jean-Frederic, David Langlais, Thiviya Jeyakumar, et al. 2024b. ‘CCDC88B Interacts with RASAL3 and ARHGEF2 and Regulates Dendritic Cell Function in Neuroinflammation and Colitis’. Communications Biology 7 (1): 77. 10.1038/s42003-023-05751-9.

O’Rourke, A. M., and M. F. Mescher. 1990. ‘T-Cell Receptor-Activated Adhesion Systems’. Current Opinion in Cell Biology 2 (5): 888–93. 10.1016/0955-0674(90)90088-V.

Qi, Yan, Ning Wang, Li-Juan Pang, et al. 2015. ‘Identification of Potential Mutations and Genomic Alterations in the Epithelial and Spindle Cell Components of Biphasic Synovial Sarcomas Using a Human Exome SNP Chip’. BMC Medical Genomics 8 (1): 69. 10.1186/s12920-015-0144-7.

Raftopoulou, Myrto, and Alan Hall. 2004. ‘Cell Migration: Rho GTPases Lead the Way’. Developmental Biology 265 (1): 23–32. 10.1016/j.ydbio.2003.06.003.

Rodriguez, R. M., C. Pitzalis, G. H. Kingsley, E. Henderson, M. J. Humphries, and G. S. Panayi. 1992. ‘T Lymphocyte Adhesion to Fibronectin (FN): A Possible Mechanism for T Cell Accumulation in the Rheumatoid Joint.’ Clinical and Experimental Immunology 89 (3): 439–45. 10.1111/j.1365-2249.1992.tb06977.x.

Saito, Suguru, Duo-Yao Cao, Aaron R. Victor, Zhenzi Peng, Hui-Ya Wu, and Derick Okwan-Duodu. 2021. ‘RASAL3 Is a Putative RasGAP Modulating Inflammatory Response by Neutrophils’. Frontiers in Immunology 12: 744300. 10.3389/fimmu.2021.744300.

Saito, Suguru, Toshihiko Kawamura, Masaya Higuchi, et al. 2015a. ‘RASAL3, a Novel Hematopoietic RasGAP Protein, Regulates the Number and Functions of NKT Cells’. European Journal of Immunology 45 (5): 1512–23. 10.1002/eji.201444977.

Saito, Suguru, Toshihiko Kawamura, Masaya Higuchi, et al. 2015b. ‘RASAL3, a Novel Hematopoietic RasGAP Protein, Regulates the Number and Functions of NKT Cells’. European Journal of Immunology 45 (5): 1512–23. 10.1002/eji.201444977.

Shin, Yoonjae, Yong Woo Kim, Hyemin Kim, et al. 2018a. ‘RASAL3 Preferentially Stimulates GTP Hydrolysis of the Rho Family Small GTPase Rac2’. Biomedical Reports 9 (3): 241–46. 10.3892/br.2018.1119.

Shin, Yoonjae, Yong Woo Kim, Hyemin Kim, et al. 2018b. ‘RASAL3 Preferentially Stimulates GTP Hydrolysis of the Rho Family Small GTPase Rac2’. Biomedical Reports 9 (3): 241–46. 10.3892/br.2018.1119.

Song, Siyang, Wenshu Cong, Shurong Zhou, et al. 2019. ‘Small GTPases: Structure, Biological Function and Its Interaction with Nanoparticles’. Asian Journal of Pharmaceutical Sciences 14 (1): 30–39. 10.1016/j.ajps.2018.06.004.

Sun, Lina, Yanhong Su, Anjun Jiao, Xin Wang, and Baojun Zhang. 2023. ‘T Cells in Health and Disease’. Signal Transduction and Targeted Therapy 8 (1): 235. 10.1038/s41392-023-01471-y.

Takeuchi, M., S. Kuramochi, N. Fusaki, et al. 1993. ‘Functional and Physical Interaction of Protein-Tyrosine Kinases Fyn and Csk in the T-Cell Signaling System’. The Journal of Biological Chemistry 268 (36): 27413–19.

Tapon, N., and A. Hall. 1997. ‘Rho, Rac and Cdc42 GTPases Regulate the Organization of the Actin Cytoskeleton’. Current Opinion in Cell Biology 9 (1): 86–92. 10.1016/s0955-0674(97)80156-1.

Toribio, M. L., J. C. Gutiérrez-Ramos, L. Pezzi, M. A. Marcos, and C. Martínez. 1989. ‘Interleukin-2-Dependent Autocrine Proliferation in T-Cell Development’. Nature 342 (6245): 82–85. 10.1038/342082a0.

Zhu, Naiqiang, and Jingyi Hou. 2020. ‘Assessing Immune Infiltration and the Tumor Microenvironment for the Diagnosis and Prognosis of Sarcoma’. Cancer Cell International 20 (1): 577. 10.1186/s12935-020-01672-3.

Zugasti, Inés, Lady Espinosa-Aroca, Klaudyna Fidyt, et al. 2025. ‘CAR-T Cell Therapy for Cancer: Current Challenges and Future Directions’. Signal Transduction and Targeted Therapy 10 (1): 210. 10.1038/s41392-025-02269-w.

