## Supplemental figures for "RASAL3 regulates RAC/CDC42 GTPases, SAPK/JNK signaling, IL-2 gene activity, and directed motility in human T cells"

#### Supplementary Figure 1

**A**

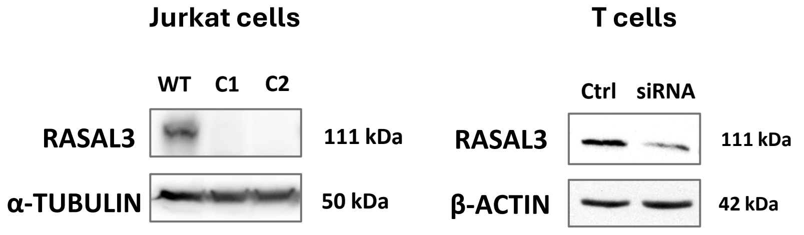


**B**

**qPCR primers sequences**

| Gene | Forward | Reverse |
|---|---|---|
| *c-Fos* | GCCTCTCTTACTACCACTCACC | TGCTGCGTTAGCATGAGTTGGC |
| *c-Jun* | CCTTGAAAGCTCAGAACTCGGAG | TGCTGCGTTAGCATGAGTTGGC |
| *Hprt* | CATTATGCTGAGGATTTGGAAAGG | CTTGAGCACACAGAGGGCTACA |
| *Gapdh* | GTCTCCTCTGACTTCAACAGCG | ACCACCCTGTTGCTGTAGCCAA |

**C**

**Single-guide RNA (electroporation sequences)**

| *rasal3* knockout |
|---|
| CCGCUUUGCUGGCUGGGGCA |
| UACCGCUGGCACACAGGGGG |
| AGGUCCGGCUUGGCGACGGU |

#### Supplementary Figure 2

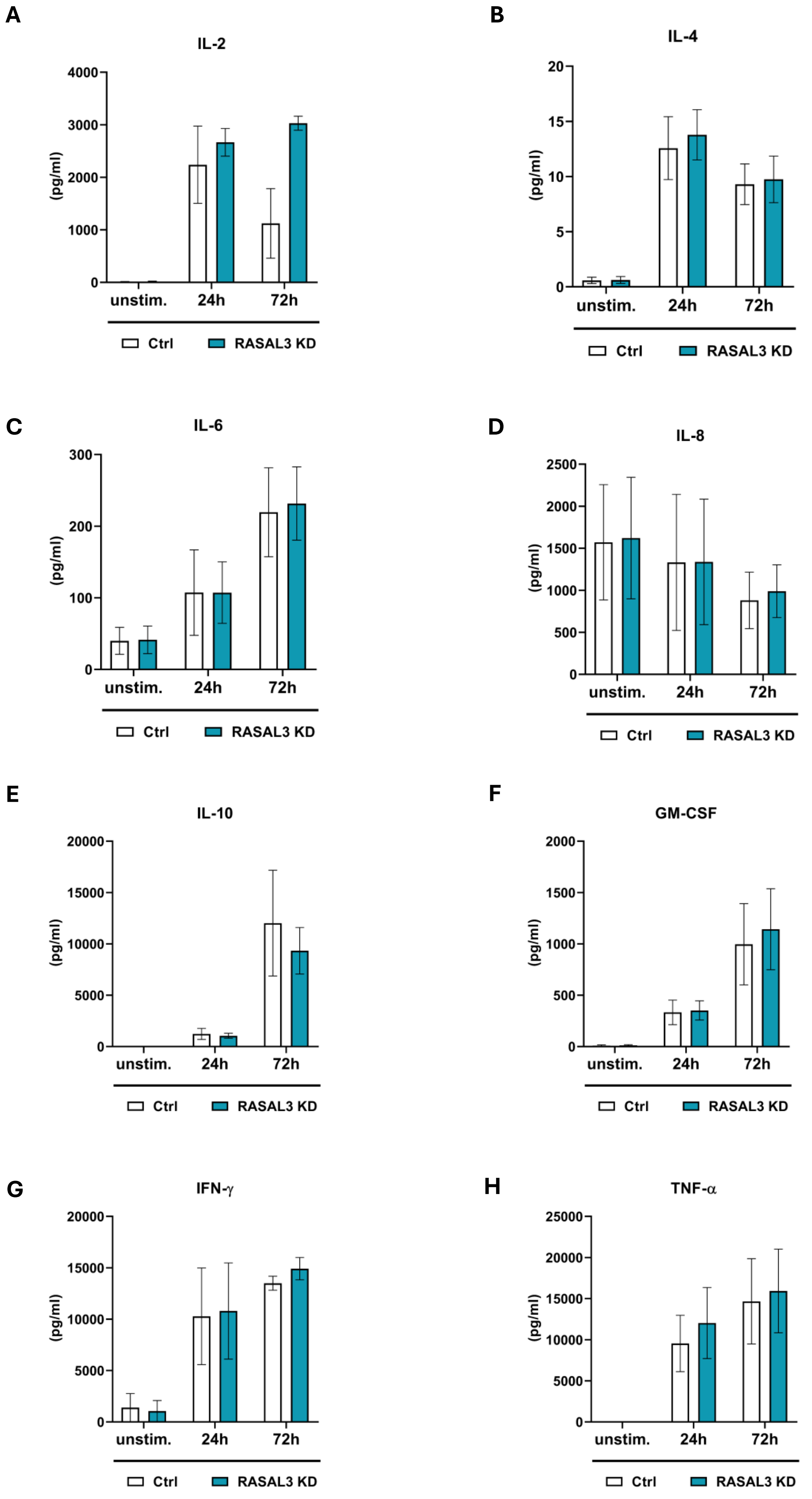

**A**

### Supplementary Figure 3

#### Western blot antibodies

| Antibody | Name | kDa | Dilution | Host/Isotype | Company |
|---|---|---|---|---|---|
| **AKT** | AKT (pan) (40D4) | 60 | 1:2000 | Mouse | CST |
| **β-ACTIN** | Monoclonal Anti-β-Actin antibody produced in mouse | 42 | 1:10000 | Mouse | SA |
| **FYN** | Anti-FYN antibody | 59 | 1:1000 | Mouse | Biosource |
| **GAPDH** | Anti-GAPDH antibody | 37 | 1:1000 | Mouse | ProteinTech |
| **GRB2** | Anti-GRB2 antibody | 25 | 1:1000 | Rabbit | Santa Cruz |
| **LAT** | Anti-LAT antibody | 38 | 1:1000 | Mouse | Exbio |
| **MAPK (ERK1/2)** | p44/42 MAPK (ERK1/2) (L34F12) | 42, 44 | 1:2000 | Mouse | CST |
| **p-AKT** | Phospho-AKT (Ser473) (D9E) | 60 | 1:2000 | Rabbit | CST |
| **p-MAPK (ERK1/2)** | Phospho-p44/42 MAPK (ERK1/2) (Thr202/Tyr204) (197G2) | 42, 44 | 1:1000 | Rabbit | CST |
| **p-P38** | Phospho-p38 MAPK (Thr180/Tyr182) (D3F9) mAb | 38 | 1:1000 | Rabbit | CST |
| **p-PLCγ1** | Anti-PLCγ1 (Y783) | 150 | 1:1000 | Rabbit | CST |
| **p-SAPK/JNK** | Phospho-SAPK/JNK (Thr183/Tyr185) (81E11) | 46, 54 | 1:1000 | Rabbit | CST |
| **p-Tyr** | Anti-phospo-Tyr (4G10) |  | 1:100 | Mouse hybridoma | Produced in house |
| **RAS** | Anti-RAS antibody | 21 | 1:200 | Mouse | TFS |
| **RAC1** | Anti-RAC1 Antibody | 21 | 1:1000 | Mouse | TFS |
| **RAC2** | RAC2 Polyclonal antibody | 21 | 1:1000 | Rabbit | P |
| **RHO** | Anti-RHO antibody | 21 | 1:666 | Rabbit | TFS |
| **RASA1** | Anti-RASA1 antibody | 120 | 1:100 | Mouse | Santa Cruz |
| **RASAL3** | RASAL3 Antibody - C-terminal region (ARP79758_P050) | 111 | 1:500 | Rabbit | ASB |
| **SRC** | Anti-Src (pY418) | 56 | 1:1000 | Rabbit | Biosource |
| **α-TUBULIN** | Anti-α-TUBULIN antibody | 50 | 1:10000 | Mouse | SA |
| **ZAP70** | Anti-ZAP70 antibody (Y319) | 70 | 1:1000 | Rabbit | CST |
